# Development of real-time polymerase chain reaction assays allowing molecular detection of *Echinococcus felidis, Echinococcus granulosus* sensu stricto, and *Echinococcus canadensis* in carnivore feces samples, animal and human hydatid cyst material from Uganda and Kenya

**DOI:** 10.1101/211920

**Authors:** Katharina Kopp

## Abstract

First evaluations on field samples, including carnivore feces, animal and human hydatid cyst material from Uganda and Kenya, showed specific amplification of two target regions of the mitochondrial genome of *Echinococcus* species according to melt and high-resolution melt curve analyses of the developed real-time PCR assays. Consecutive sequencing of PCR products revealed that, apart from *Echinococcus felidis*, sequences of two other tapeworm species, *Echinococcus granulosus* sensu stricto and *Echinococcus canadensis*, which are also endemic in East Africa, were detected by the developed real-time PCR assays.

## 0. Introduction

The aim of the present study was to develop a rapid, sensitive, and safe method for molecular detection of *Echinococcus felidis*, and possibly, related tapeworm species. We first investigated nucleic acid extraction methods suitable for isolating templates directly from fecal samples of carnivores and hydatid cyst material of intermediate hosts in the field without tedious laboratory based enrichment methods, such as digestion or flotation methods. Then, we developed real-time PCR assays which target two different mitochondrial genome regions of *E. felidis*.

*Echinococcus granulosus* sensu stricto (s.s.) and *Echinococcus canadensis* are known to be zoonotic parasites (Alvarez Rojas et al., 2014). In East Africa, both tapeworm species pose a public health burden, mainly on (semi-)nomadic pastoralist communities in remote areas (Wachira et al., 1993, Dinkel et al., 2004, Magambo et al., 2006, Hüttner et al., 2008, 2009, Hüttner and Romig, 2009, Casulli et al., 2010, Omer et al., 2010, Romig et al., 2011, Addy et al., 2012, Mutwiri et al., 2013, Mbaya et al., 2014, Addy et al., 2017), as they can cause cystic echinococcosis (CE), a severe chronic disease in human patients. Cystic echinococcosis has been recognized as a neglected zoonotic disease (NZD) and a neglected tropical disease (NTD) by the World Health Organization (WHO) (http://www.who.int/entity/neglected_diseases/en/, as accessed on 15 Sept 2017).

However, little is known about the geographical distribution, epidemiology, and ecology of the “lion strain” tapeworm *E. felidis.* Known definitive hosts are African lions (*Panthera leo*) and spotted hyenas (*Crocuta crocuta*) (Hüttner et al., 2008, 2009, Kagendo et al., 2014). One warthog and six hippopotami have been identified as intermediate hosts to date (Hüttner et al., 2009, Halajian et al., 2017). The only countries in which the presence of *E. felidis* has been confirmed are Uganda, Kenya, and South Africa (Hüttner et al,. 2008, Hüttner and Romig, 2009, Hüttner et al., 2009, Kagendo et al., 2014, Halajian et al., 2017). While the complete mitochondrial genome of *E. felidis* has been sequenced from an isolated taeniid egg (Nakao et al, 2013), the whole nuclear genome has not been sequenced. Until now, *E. felidis* has not been identified as causative agent of any human CE case (Romig et al., 2017). Fast, sensitive, and fieldable molecular detection methods, such as the developed real-time PCR assays, could provide useful tools in epidemiological and ecological large-scale studies aiming to close this gap in knowledge on *E. felidis*.

## 1. Materials and Methods

### 1.1 Sample collection and processing

Two sample batches were collected, preserved, and extracted as listed in Table 1 and 2. The first batch of samples (Table 1) originates from wildlife inhabiting Queen Elisabeth National Park (QENP) in Western Uganda. The second batch of samples came from Kenya, and included materials from two human cystic echinococcosis (CE) patients, a wild carnivore (F3, unknown species), and a domestic dog (Table 2). After collection, all samples were immediately mixed with an equal volume (1:1) of 80% (vol/vol) ethanol and stored at 4-8°C temperature. However, some samples were transported for up to three days under ambient tropical temperatures before being stored at 4-8°C again.

**Table 1:** Samples from Uganda. See detailed description of nucleic acid extraction methods GL and TLN under respective paragraph in chapter Materials and Methods. Unknown (u), male (Ma).

| Sample name | Sample type | Host species | Gender | Location | Preservation liquid | Remarks | Sampling date/ID | Extraction date | Extraction method |
| --- | --- | --- | --- | --- | --- | --- | --- | --- | --- |
| GL2 | feces | African lion | u | QENP | 80% ethanol | #06 yellow cap Falcon 50 ml | u | 21 April 2017 | GL |
| TLN18 | feces | African lion | Ma | QENP | 80% ethanol | Kadogo | 20170330 | 26 April 2017 | TLN |
| TLN5 | "skin" chest tissue | giant forest hog | u | u | 80% ethanol | Kass | 20170217 | 26 April 2017 | TLN |

**Table 2.** Samples from Kenya. See detailed description of nucleic acid extraction methods ZM and pFl under respective paragraph in chapter Materials and Methods. Unknown (u), none (n).

| Sample name | Sample type | Host species | Gender | Location | Preservation liquid | Remarks | Sampling date/ID | Extraction date | Extraction method |
| --- | --- | --- | --- | --- | --- | --- | --- | --- | --- |
| M5 | meta-cestode | human | u | Kenya | 80% ethanol | n | u | 23 April 2017 | ZM |
| M6 | meta-cestode | human | u | Kenya | 80% ethanol | n | u | 24 April 2017 | ZM |
| F3 | feces | unknown wild carnivore | u | Kenya | n.a., dried | n | u | 24 April 2017 | ZM |
| F4-df | feces | domestic dog | u | Kenya | 80% ethanol | n | u | 23 April 2017 | ZM |
| F4-pFI |  |  |  |  |  |  |  | 24 April 2017 | pFI + ZM |

### 1.2 Nucleic Acid Extraction Methods

During this proof of principle study several nucleic acid extraction methods were available to us. Therefore, various samples were extracted by different methods. However, all sample processing protocols employed a mechanical homogenization step using a bead lysis matrix and the TerraLyzer™ device (S6022, Zymo Research Corp., Irvine, CA, USA). After elution, all samples were stored at −20°C before using them as templates in the developed PCR assays.

**GL**: Total DNA/RNA was extracted and eluted in 100 µl nuclease-free water from one fecal-ethanol mixture sample (GL2) by using the Quick-RNA MiniPrep kit (R1054, Zymo Research Corp., Irvine, CA, USA). The protocol of the manufacturer was slightly modified. Instead of the recommended ZR Bashing Beads™, 0.7 ml dry volume of locally sourced glass seed beads ("Gazaland mall", Kampala, Uganda), which are commonly used for artisan jewelry in East Africa, was employed as lysis matrix in a 2.0 ml micro tube (2.0 ml SC Micro Tube PCR-PT, 72.694.700, Sarstedt AG & Co., Nürmbrecht, Germany). Similar commercial beads are available internationally, e.g. TOHO Demi Round 11/0 2.2 mm Seed Beads (#TN-11-PF21F, Starman Inc., Sequim, WA, USA). After adding 300 µl of the kit's proprietary RNA

Lysis Buffer and 200 µl of sample, the mixture was homogenized mechanically by using the TerraLyzer™ device for 1 min. The consecutive steps followed the manufacturer's protocol but did not apply a DNase digestion step. Total DNA/RNA was eluted with 100 µl nuclease-free water.

**TLN**: The original content of a lysis tube (R1105, Zymo Research Corp., Irvine, CA, USA) was equally distributed to two 2.0 ml micro tubes (2.0 ml SC Micro Tube PCR-PT, 72.694.700, Sarstedt AG & Co., Nürmbrecht, Germany), so that each resulting micro tube contained 0.5 ml of DNA/RNA Shield™ and 0.35 ml dry volume 2.0 mm ZR Bashing Beads™. After reception of samples from the field (QENP), where they had been stored at 4-8°C in 80% (vol/vol) ethanol, 300 µl of each sample-ethanol mixture (GL2 and TLN18) and 300 mg of tissue (TLN5) were transferred to a prepared micro tube (see above). Each sample in a micro tube was thoroughly homogenized by using the TerraLyzer™ device (Zymo Research Corp., Irvine, CA, USA) for 1 min and stored at 4-8°C. Fourteen days after this pre-treatment, a volume of 100 µl of the homogenized sample solution was mixed with 300 µl TRI Reagent™ (R2050-1-200, Zymo Research Corp., Irvine, CA, USA) and extracted by using the Directzol kit™ (Zymo Research Corp., Irvine, CA, USA) without DNase digest according to the manufacturer's protocol. Total DNA/RNA was eluted with 100 µl nuclease-free water.

**ZM**: The samples (M5, M6, and F4-df) which were processed by using this method were stored in 80% (vol/vol) ethanol (with exception of dried fecal sample F3) and shipped at ambient temperature (25-40°C) for three days. One ml of dried fecal sample F3 was mixed with 9 ml 80% (vol/vol) ethanol before further processing. For DNA extraction from samples the ZymoBIOMICS™ DNA Miniprep kit (Zymo Research Corp., Irvine, CA, USA) was used. Each 2.0 ml lysis tube of this kit contained 0.7 ml dry volume of a mixture of 0.1 & 0.5 mm ZR Bashing Beads™. A volume of 750 µl proprietary ZymoBIOMICS™ Lysis Solution was added to each tube, as well as 200 mg of human hydatid cyst material (M5 and M6) and 200 µl of fecal sample-ethanol mixture (F3 and F4), respectively. Sample homogenization was achieved by 1 min rigorous mechanical shaking using the TerraLyzer™ device. The following DNA extraction steps were undertaken as recommended by the manufacturer. Total DNA was eluted in 100 µl of nuclease-free water.

**pFl**: This method was only applied to an aliquot of sample F4 (F4-pFl) and was modified from the protocol described in Széll et al., 2014. Briefly, a volume of 1 ml of feces-ethanol sample was mixed with 3.2 ml tap water, strained through a sieve (0.6 mm mesh), transferred to two 16 ml tubes, and centrifuged at 2000 g for 10 min. The supernatant was discarded and the sediment was re-suspended in 10 ml modified Breza solution. After another centrifugation at 2000 g for 15 min the surface of the flotation fluid was touched with the end of a glass rod and transferred onto a microscope slide. The latter was repeated four times. If taeniid eggs were found by microscopic inspection, the surface area of the microscope slide where the sample's drops had been transferred as well as the bottom part of the cover slide were rinsed with a total volume of 200 µl of nuclease-free water. Further nucleic extraction steps from the entire collected volume of the resulting eggs-in-water suspension followed the procedure as described under method "ZM" (see above). Total DNA was eluted in 100 µl of nuclease-free water.

### 1.3 Real-time (RT-)PCR assays

Two polymerase chain reaction (PCR) assays were developed. For each PCR assay a pair of primers was designed targeting different genes. The assays can be used as end-point (conventional) PCR run. However, the aim of this study was the proof of principle of molecular detection of *Echinococcus* sequences by using real-time PCR. Therefore, the analysis of cycling data and of simple and high-resolution melt curve were established. We also performed the developed PCR assays as one-step reverse transcription (RT) real-time PCR runs on total RNA and DNA extracts.

#### 1.3.1 Primer design

Primers were designed in order to target two regions of the *E. felidis* mitochondrial DNA, complete genome, sample code EfelUganda (GenBank entry AB732958.1). We chose two mitochondrial gene targets. The first gene cytochrome c oxidase subunit 1 (cox1) is located at position 9066-10673 of the mitochondrial genome. The second gene NADH dehydrogenase subunit 5 (nad5) is located upstream at position 453-2024. Primer pairs were selected by using the tool Primer3web version 4.0.0 (http://primer3.ut.ee/, Untergasser et al., 2012, Koressaar and Remm, 2007). Primer specificity was tested by using the NCBI Primer-BLAST tool (https://www.ncbi.nlm.nih.gov/tools/primer-blast/, Ye et al., 2012) against the non-redundant GenBank database "nr".

#### 1.3.2 Reaction set-up

In order to achieve equal reagent concentrations, all real-time PCR and RT-PCR reactions were set up as bulk master mixture with the respective components of the GoTaq™ 1-Step RT-qPCR System for Dye-Based Detection (A6020, Promega Corp., Madison, WI, USA). Each real-time (RT-)PCR assay was performed in a final reaction volume of 20 µl, containing 10 µl GoTaq™ qPCR Master Mix, 2X, 4 pmol of each primer (forward and reverse primer of set Ec-1 or set Ec-2), and 5.2 µl nuclease-free water (PCR assay), or 4.8 µl nuclease-free water and 0.4 µl GoScript™ RT Mix for 1-Step RT-qPCR, 50X (RT-PCR assay). To each of these mixture volumes of 16 µl either 4 µl of total nucleic acid extract from a sample was added as template or 4 µl of nuclease-free water in no-template controls. For each specimen the reaction was set up in duplicate, triplicate, or sextuplicate. All real-time PCR and RT-PCR assays were run on the Magnetic Induction Cycler micPCR™ system (Bio Molecular Systems, Potts Point, NSW, Australia) using tubes (REF 60653, Bio Molecular Systems) with a maximum volume of 25 µl, which had been preloaded with silicone oil and were closed by respective caps.

#### 1.3.3 Cycling conditions

Cycling profiles were set up and controlled as well as results monitored and analyzed by using the micPCR™ software suite version 1.8.0 (copyright © 2014-2015, Bio Molecular Systems, Potts Point, NSW, Australia). As the hot-start DNA polymerase of the used GoTaq™ qPCR Master Mix (Promega Corp., Madison, WI, USA) is compatible with fast cycling conditions according to the manufacturer, Fast TAQ (v3) temperature control mode was chosen for all cycling profiles. All real-time RT-PCR assays were run as one-step reactions without opening of reaction tubes. Reverse transcription of RNA, potentially present in total DNA/RNA extracts, was attempted by holding the reaction at 50°C for 15 min.

The following cycling steps for amplification and real-time detection of intercalating dye fluorescence in real-time RT-PCR assays also applied for the real-time PCR assays: initial denaturation of the template, inactivation of the reverse transcriptase (in RT-PCR assays only), and activation of the hot-start DNA polymerase at 95°C for 10 min; 10 pre-cycles with 95°C for 10 s, 60°C for 30 s with a temperature decrease of 0.8°C per cycle, and 72°C for 30 s; and 30 cycles with 95°C for 10 s, 52°C for 30 s, and 72°C for 30 s during which the fluorescence was read. Since Promega's proprietary BRYT Green™ dye has spectral properties similar to those of SYBR™ Green I dye, the latter chemistry was selected for the micPCR system's parameters.

#### 1.3.4 Melting conditions

As intercalating dyes bind non-specifically into PCR products (Morrison et al., 1998), a melting of PCR products was performed after PCR cycling in order to identify the correct amplicon by its specific melting characteristics. Melting included 95°C for 5 s, 65 °C for 15 s, and heating to 95°C at a rate of 0.1°C/s with continuous reading of fluorescence.

#### 1.3.5 Cycling data analysis

The quantification cycle (*C_q_*) value of each real-time (RT)-PCR reaction, as defined by MIQE guidelines (Bustin et al., 2009), was calculated by using the following parameters of the micPCR version 1.8.0. software. The option "Dynamic" (see micPCR™ User Manual 1.8.0, copyright © 2016, Bio Molecular Systems, Potts Point, NSW, Australia, for details) was chosen as baseline correction method, the threshold was automatically set by the analysis software, and a fluorescence cutoff level of 5% was selected. By default, baseline-corrected curves were plot as fluorescence (y-axis) against cycle number (x-axis) for single reactions, in logarithmic scale. As each reaction containing sample template was run in duplicate, triplicate, or sextuplicate, average *C_q_* values were calculated.

#### 1.3.6 Melt curve analysis

The peak dissociation temperature (*Tm*) of each PCR product was determined from the local maximum of the first derivative curve plotted as -*dF/dT* (y-axis) against temperature (°C, x-axis). The melt curve threshold was set to a common value of 0.064, with two exceptions in which the amplitude of the *Tm* peak was found to be low (a threshold of 0.060, see Figure 3 and a threshold of 0.021, see Figure 5).

#### 1.3.7 High-resolution melt curve analysis

The high-resolution melt (HRM) curve displays the normalized fluorescence (y-axis) plotted against temperature (°C, x-axis). For all (RT-)PCR assays the following two normalization regions were chosen. The first region was set to 68.0-71.0°C, the second region to 92.0-93.5°C, and the confidence limit to 90%.

### 1.4 Sequencing of real-time PCR products

Real-time PCR products which were determined as specific amplification according to melt curve analysis and HRM analysis were pooled for duplicate, triplicate, or sextuplicate reactions and purified by using PCR and Gelextraction Mini Prep Kit (S5380.0050, Genaxxon bioscience GmbH, Ulm, Germany). Forward (using primer Ec-1-FW or Ec-2-FW) and reverse (using primer Ec-1-REV or Ec-2-REV) strands of each of these cleaned amplicons were sequenced using the Sanger method (Sanger et al., 1977) by Macrogen Europe Inc., Amsterdam, the Netherlands.

### 1.5 Bioinformatics data analysis of real-time PCR product sequences

For each sequenced PCR product the reported forward and reverse strands were visualized, aligned and merged to one amplicon sequence by using Jalview (version 2, last updated: 04 June 2014, Waterhouse et al., 2009). A search over the entire "nucleotide collection (nr/nt)" database by using the "blastn suite" https://blast.ncbi.nlm.nih.gov/Blast.cgi?PROGRAM=blastn) of the BLAST algorithm (Altschul et al., 1990) was conducted for the resulting merged amplicon sequences. No low complexity region filter and no mask were used.

For each merged PCR product the sequence of the GenBank entry which matched best according to the BLAST search was recorded. Two multiple sequence alignments (MSA) were constructed of merged PCR products and respective best BLAST match sequences by using Clustal Omega (http://ww.ebi.ac.uk/Tools/msa/clustalo/, Sievers et al., 2011). One MSA represented sequences associated with primer pair Ec-1, the other MSA contained sequences with regard to primer pair Ec-2.

Both MSAs were further analyzed by using Jalview. A pairwise alignment of every single sequence of a MSA against each other sequence was conducted and the resulting percentage of identity (PID in %) for each pair recorded. The resulting PID matrices were analyzed regarding grouping of PCR products and *Echinococcus* species of similar GenBank sequences.

Apart from the best BLAST matches, further non-redundant corresponding regions of other *Echinococcus* species were added to each of the two MSAs. From these extended MSAs two phylogenetic trees based on average distance of PID were calculated using Jalview. The unweighted pair-group method using arithmetic averages (UPGMA) (Sneath and Sokal, 1973, Michener and Sokal, 1957) was employed as agglomerative clustering algorithm. Clusters were iteratively formed and extended by finding a non-member sequence with the lowest average dissimilarity over the cluster members.

## 2. Results and Discussion

Samples in which nucleotide sequences of the genus *Echinococcus* were detected by the developed real-time (RT-)PCR assays and confirmed by Sanger sequencing of the PCR products showed a distinct species separation. All samples from Uganda were identified as containing DNA of *Echinococcus felidis*, while the samples from Kenya belonged to genotype G1 and G3 of *Echinococcus granulosus* sensu stricto (s.s.) and genotype G6 of *Echinococcus canadensis*. Due to reagent availability also different nucleic acid extraction methods were applied for samples from these two neighboring East African countries. A common total DNA extraction method (ZM, see chapter materials and methods) was applied to all Kenyan samples. Two different methods (GL, TLN, see chapter materials and methods) were used for total DNA and RNA extraction in all Ugandan samples, though.

### 2.1 Primer pairs detected two gene targets of three *Echinococcus* species

Primer design resulted in two primer pairs (Table 3). The first pair Ec-1-FW and Ec-1-REV targets a 372 bp long region ranging from the downstream stretch of the gene cytochrome c oxidase subunit 1 (cox1), which overlaps with the region of the gene tnrT at position 10668-10746 (for GenBank entry AB732958.1), to an intergenic region. The second primer pair Ec-2-FW and Ec-2-REV targets a 309 bp long region located within the downstream third of the gene NADH dehydrogenase subunit 5 (nad5).

**Table 3:**
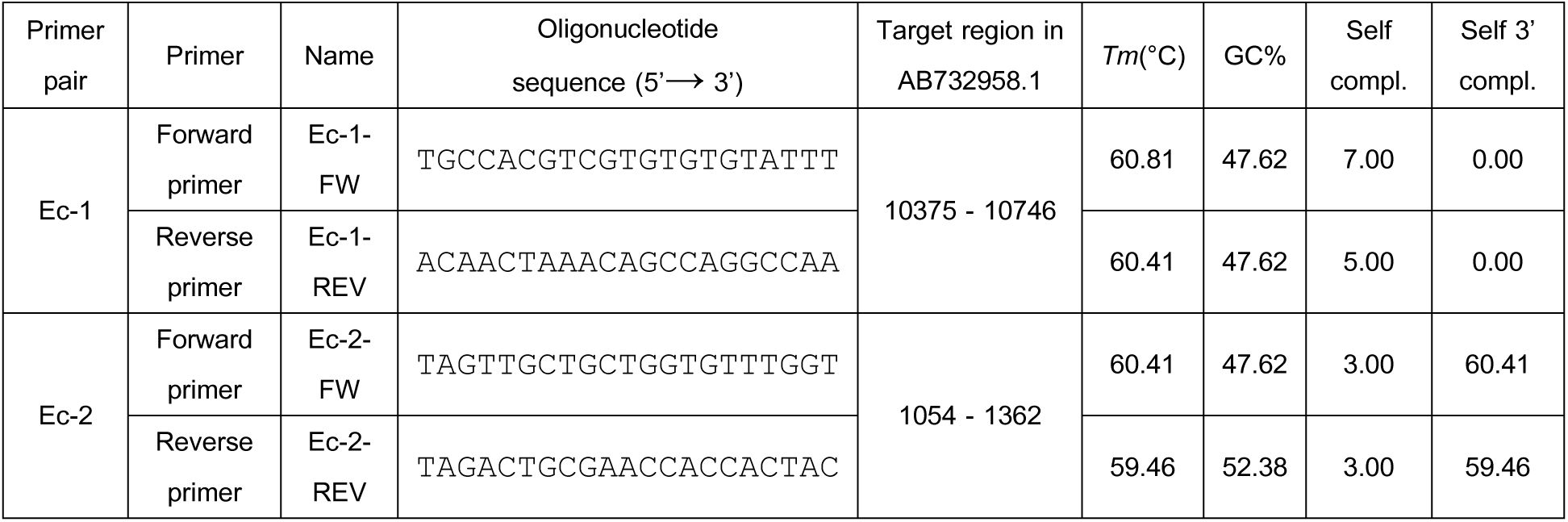
Primer pairs Ec-1 and Ec-2 for real-time PCR assays targeting cox1/trnT and nad5 gene regions in *E. felidis*. Annealing temperature *T_m_* (°C), percentage of GC content (GC%), self-complementarity of primer sequence (Self compl.) and of primer sequence 3'-terminal end (Self 3' compl.).

The primer sets Ec-1 and Ec-2 had been originally designed to specifically target two regions within the mitochondrial genome of *E. felidis* only. However, when the PCR assays were used in several field samples, we observed melt curve peaks and high resolution melting curve profiles which are typical for specific amplification, although consecutive sequencing of the respective PCR products found that the DNA templates belonged to *E. granulosus* s.s. or *E. canadensis*. Some level of unspecific amplification is known to occur especially for dye based real-time PCR assays in contrast to assays which employ hydrolysis probes. This applies especially, if single nucleotide mismatches are present only at the 5' terminus of a primer with the template. While even several mismatches between template and primer are well tolerated at the 5' terminal end of the primer, mismatches at the 3' terminal tend to hamper primer annealing to the template and extension of the respective PCR product.

When PCR products of assays using the primer pair Ec-1 were sequenced, BLAST searches gave best matches for sequences of *E. felidis* for samples from Uganda and *E. granulosus* s.s. (genotype G1 and G3) for samples from Kenya. This is reflected in the alignments of primers Ec-1-FW and Ec-1-REV with best matching GenBank entry sequences (Table 4). As intended by original primer design forward and reverse primers of pair Ec-1 are identical with the target region of the mitochondrial genome of *E. felidis*. While Ec-1-FW matches perfectly with the respective target region sequence of *E. canadensis*, a total of five mismatches between Ec-1-REV and the target sequence of this species occur, with two mismatches located at the 3' terminal third of the primer. In line with this alignment, none of the sequenced PCR products of primer pair Ec-1 was determined to belong to the species *E. canadensis*. Two sequenced PCR products of assays using primer pair Ec-1 were found by BLAST searches to match best to sequences of *E. granulosus* s.s. (genotype G1 and G3), while two mismatches with the sequence of primer Ec-1-FW and three mismatches with the sequence of primer Ec-1-REV were found for the alignments with the closest GenBank entries KY766892.1 and KY766891.1. However, only one mismatch is located in the 3'-terminal third of each of the two primers.

When PCR products of assays using the primer pair Ec-2 were sequenced, BLAST searches gave best matches for sequences of *E. felidis* for samples from Uganda. For samples from Kenya best matches were found for sequences of *E. granulosus* s.s. (genotype G1 and G3) and of *E. canadensis* genotype G6. This is reflected in the alignments of primers Ec-2-FW and Ec-2-REV with best matching GenBank entry sequences (Table 5). As found for primer pair Ec-1, and intended by original primer design, forward and reverse primers of pair Ec-2 are identical with the target region of the mitochondrial genome of *E. felidis*. This applies also for primer Ec-2-FW with respect to the respective target region sequence of *E. canadensis* genotype G6, while a single mismatch was found within the 5' terminal third of primer Ec-2 REV. In accordance with this alignment, sequencing of two PCR products identified them as belonging to *E. canadensis* genotype G6. This sequence region of *E. canadensis* genotype G6 is identical with the respective region of GenBank entry KC756275.1, while it had been assigned to *E. granulosus* "genotype G6" in this GenBank record. As discussed later, due to recent phylogenetic studies (Addy et al., 2017), this entry should rather be classified as also belonging to the species *E. canadensis*. Two sequenced PCR products of assays using primer pair Ec-2 were found by BLAST searches to match best to sequences of *E. granulosus* s.s. (genotype G1 and G3). One mismatch within the 5-terminal third of the primer Ec-2-FW was found for the GenBank entries representing these two genotypes of *E. granulosus* s.s. Two and three mismatches with the sequence of primer Ec-2-REV were found for the alignments with the closest GenBank entries KY766892.1 and KY766891.1. However, as determined for the reverse primer of pair Ec-1, only one mismatch is located in the 3'-terminal third of the primer Ec-2-REV.

**Table 4:**
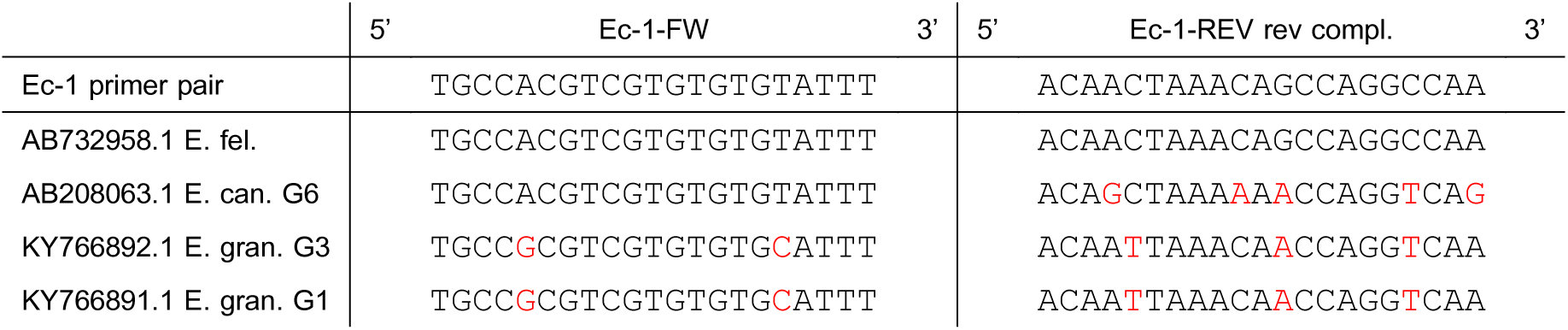
Alignment of forward and reverse primers of pair Ec-1 to GenBank entries corresponding to the target region (cox1) in the mitochondrial genome of *E. felidis*, mismatches with primer regions marked in red.

**Table 5:**
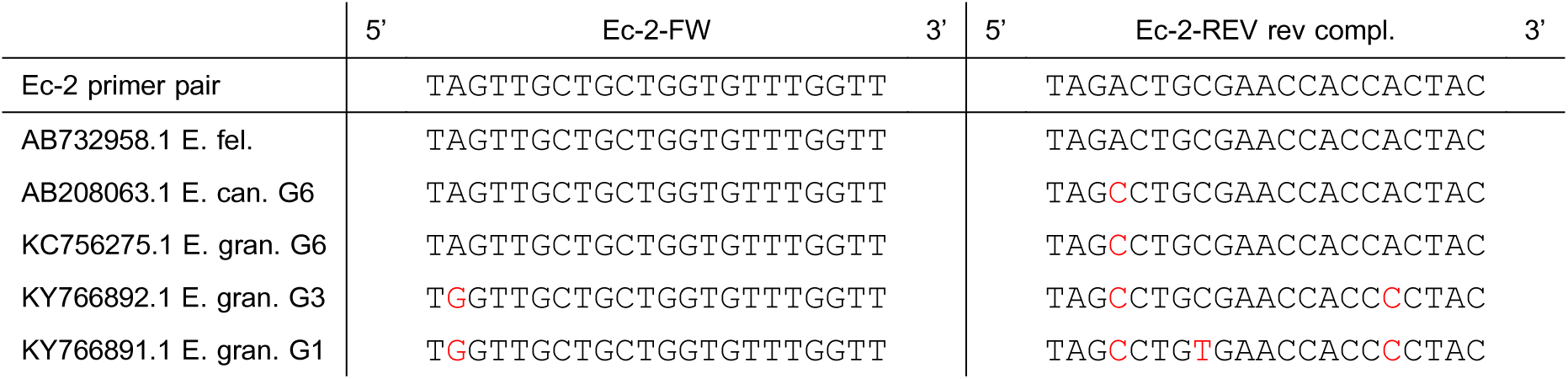
Alignment of forward and reverse primers of pair Ec-2 to GenBank entries corresponding to the target region (nad5) in the mitochondrial genome of *E. felidis*, mismatches with primer regions marked in red.

### 2.2 Samples from Uganda tested positive in real-time PCR assays and were confirmed by Sanger sequencing of the PCR products

Real-time PCR of two samples of lion feces collected from different animals in Queen Elizabeth National Park (QENP), Western Uganda and extracted with different nucleic acid extraction methods, resulted in detection of two different target genes, NADH dehydrogenase subunit 5 (nad5) and cytochrome c oxidase subunit 1 (cox1), of the genus *Echinococcus*. A chest tissue sample of a giant forest hog, also collected in QENP, was determined as positive in the developed PCR assay targeting the *Echinococcus* gene nad5. All respective PCR products showed specific amplification by melt curve and high resolution melt (HRM) curve analysis and were identified by Sanger sequencing as belonging to *E. felidis*. Assay conditions (used primer pair, previous reverse transcription or not), label of the resulting PCR product, melt curve analysis peak (*T_m_*), HRM curve analysis result, quantification cycle (*C_q_*), and the percentage of sequence identity of each amplicon with the respective sequence of *E. felidis* are listed in Table 6.

**Table 6:** Conditions and results of real-time (RT-)PCR assays of samples from Uganda which tested positive and were confirmed by Sanger sequencing of the PCR products. Sample name (GL2, TLN18, and TLN5, see details in Table 1 in Materials and Methods), primer pair (Ec-1- FW/-REV or Ec-2-FW/-REV), sequence ID of PCR product, reverse transcription (RT) prior to PCR performed (+) or not (-), specific high resolution melting (HRM) curve profile determined (+), average temperature (°C) of melt curve analysis peak (*T*_*m*_), average quantification cycle (*C*_*q*_), and percentage of sequence identity (% id seq) of PCR product with *E. felidis*.

| Sample | Primer set | Sequence ID | RT | HRM | $T_m$ (°C) | $C_q$ | % id seq<br>E. fel. |
| --- | --- | --- | --- | --- | --- | --- | --- |
| GL2 | Ec-2 | B1 | - | + | 80.19 | 31.61 | 100.00 |
|  |  | B8 | + | + | 80.84 | 32.85 | 98.71 |
|  | Ec-1 | B3 | - | + | 79.80 | 32.55 | 100.00 |
| TLN18 | Ec-2 | B2 | - | + | 80.34 | 33.56 | 99.35 |
|  |  | B5 | + | + | 80.17 | 32.85 | 96.76 |
|  | Ec-1 | B4 | - | + | 79.72 | 34.44 | 99.39 |
| TLN5 | Ec-2 | B9 | + | + | 80.74 | 36.22 | 97.68 |

The two lion feces samples GL2 and TLN18 showed specific amplification in two different assay conditions (RT +, RT -) each using primer set Ec-2. One assay set-up was potentially capable of performing reverse transcription of template RNA before amplification of DNA (RT +), the other could only amplify target DNA (RT -). In the first instance, the reason for attempting reverse transcription of template RNA was that some of the samples were also tested for the presence of RNA viruses, such as rabies virus or paramyxoviruses, in addition to determining the presence of *Echinococcus* DNA in one real-time (RT)-PCR run together. While different primer pairs were added in the reaction mixture testing for virus RNA and *Echinococcus* DNA, we otherwise used the same master mix (GoTaq™ qPCR Master Mix, 2X, Promega Corp., Madison, WI, USA) and run profile (constant temperature hold for reverse transcription before polymerase chain reaction cycles). The chest tissue sample TLN5 of a giant forest hog showed specific amplification in a real-time PCR assay (RT-) using primer set Ec-2. In real-time PCR assays (RT-) in which primer pair Ec-1 was used the lion feces samples GL2 and TLN18 also displayed melt curve and HRM curve analysis results which were considered specific for correct target amplification.

The respective graphs display the specific melt curve analysis peak *T*_*m*_ (Figure 1, 3, 5, and 7) and the specific HRM curve profile (2, 4, 6, and 8) in comparison with the *T*_*m*_ and HRM curve profile of the no-template controls (NTCs).

The importance of running each sample at least in duplicate can be seen in the graphs displaying melt curve and HRM results for the lion feces sample TLN18 (Figure 5 and 6 for primer pair Ec-2 and Figure 7 and 8 for primer pair Ec-1) and the giant forest hog tissue sample TLN 5 (Figure 3 and 4). As indicated by relatively high *C*_*q*_ values (32.85 - 36.22) in Table 6, the *Echinococcus* DNA amount extracted from these samples was quite low. Nevertheless, Sanger sequencing of PCR products using these samples confirmed specific amplification of target regions of the genome of *E. felidis*. *T*_*m*_ values in melt curve analyses and shapes of HRM curves tend to become ambiguous for samples containing low amounts of DNA templates close to the limit of detection (LOD) of an assay. Therefore, it is strongly recommended, if resources allow, examining each sample in triplicate. Otherwise the risk of false negatives arises.

**Figure 1:**
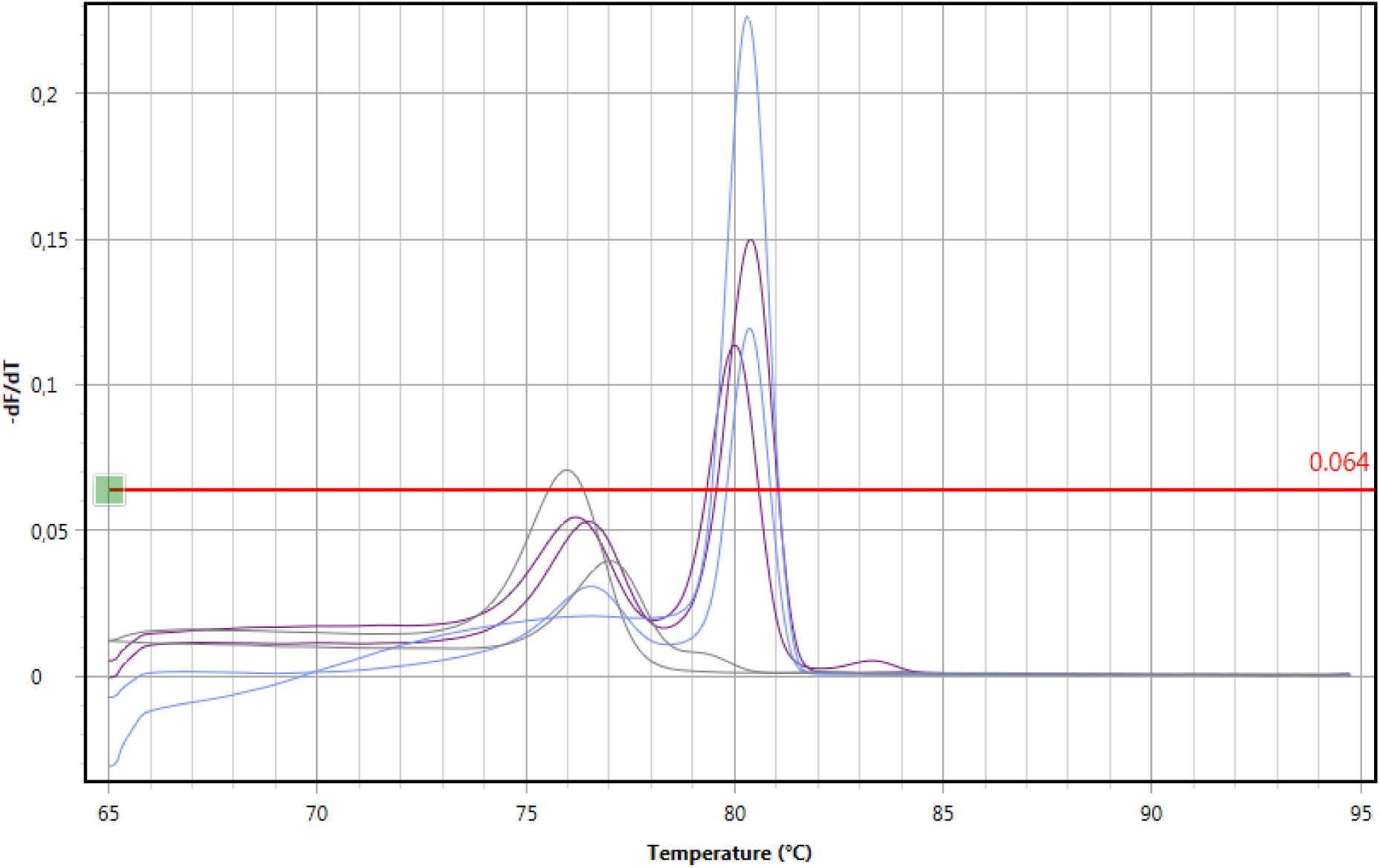
Melt curve of polymerase chain reaction of sample GL2 (dark purple), TLN18 (light blue), no template control (grey) without preceding reverse transcription using primer pair Ec-2, each reaction in duplicate.

**Figure 2:**
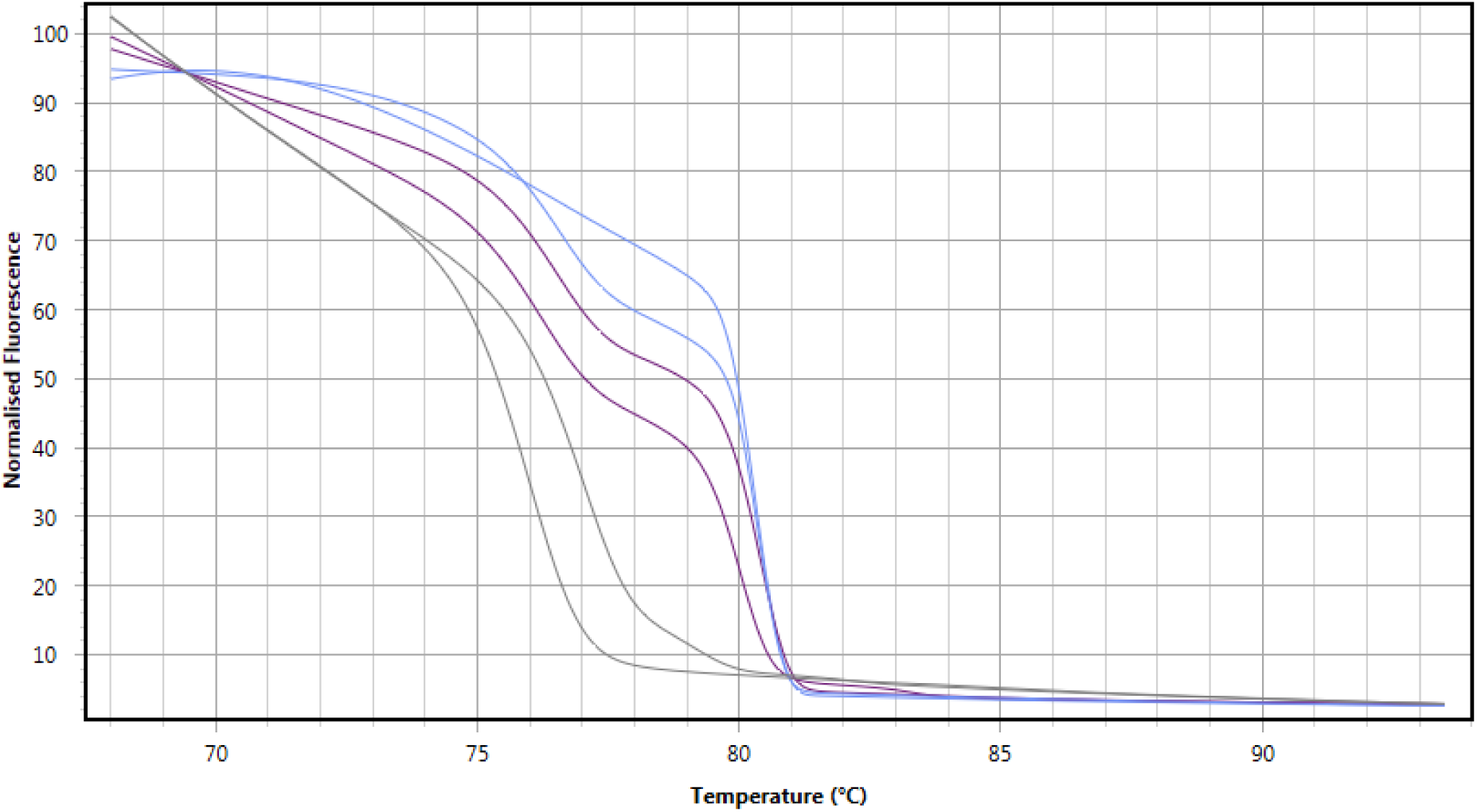
High resolution melt curve of polymerase chain reaction of sample GL2 (dark purple), TLN18 (light blue), no template control (grey) without preceding reverse transcription using primer pair Ec-2, each reaction in duplicate.

**Figure 3:**
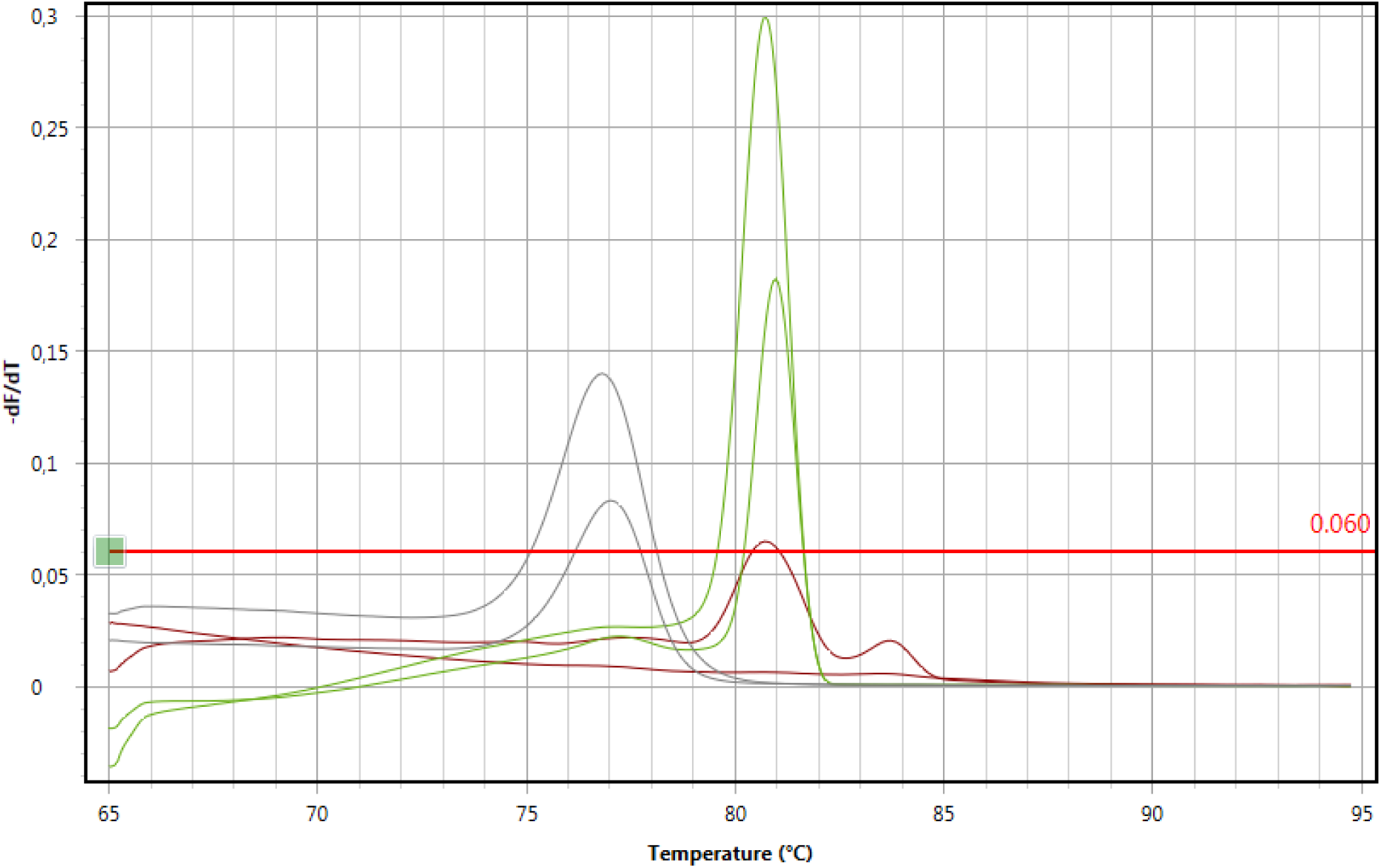
Melt curve of polymerase chain reaction of sample GL2 (green), TLN5 (brown), no template control (grey) with preceding reverse transcription using primer pair Ec-2, each reaction in duplicate.

**Figure 4:**
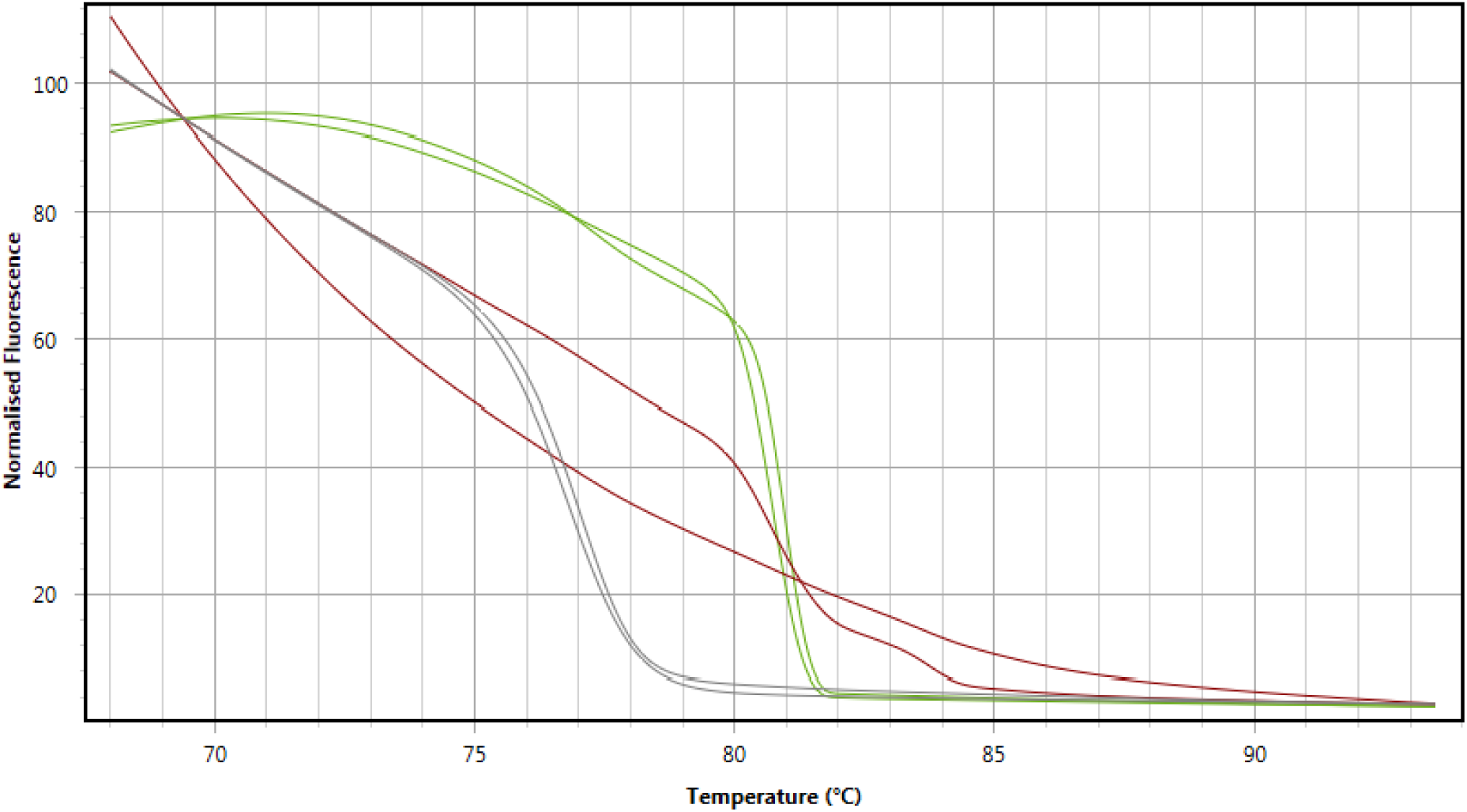
High resolution melt curve of polymerase chain reaction of sample GL2 (green), TLN5 (brown), no template control (grey) with preceding reverse transcription using primer pair Ec-2, each reaction in duplicate.

**Figure 5:**
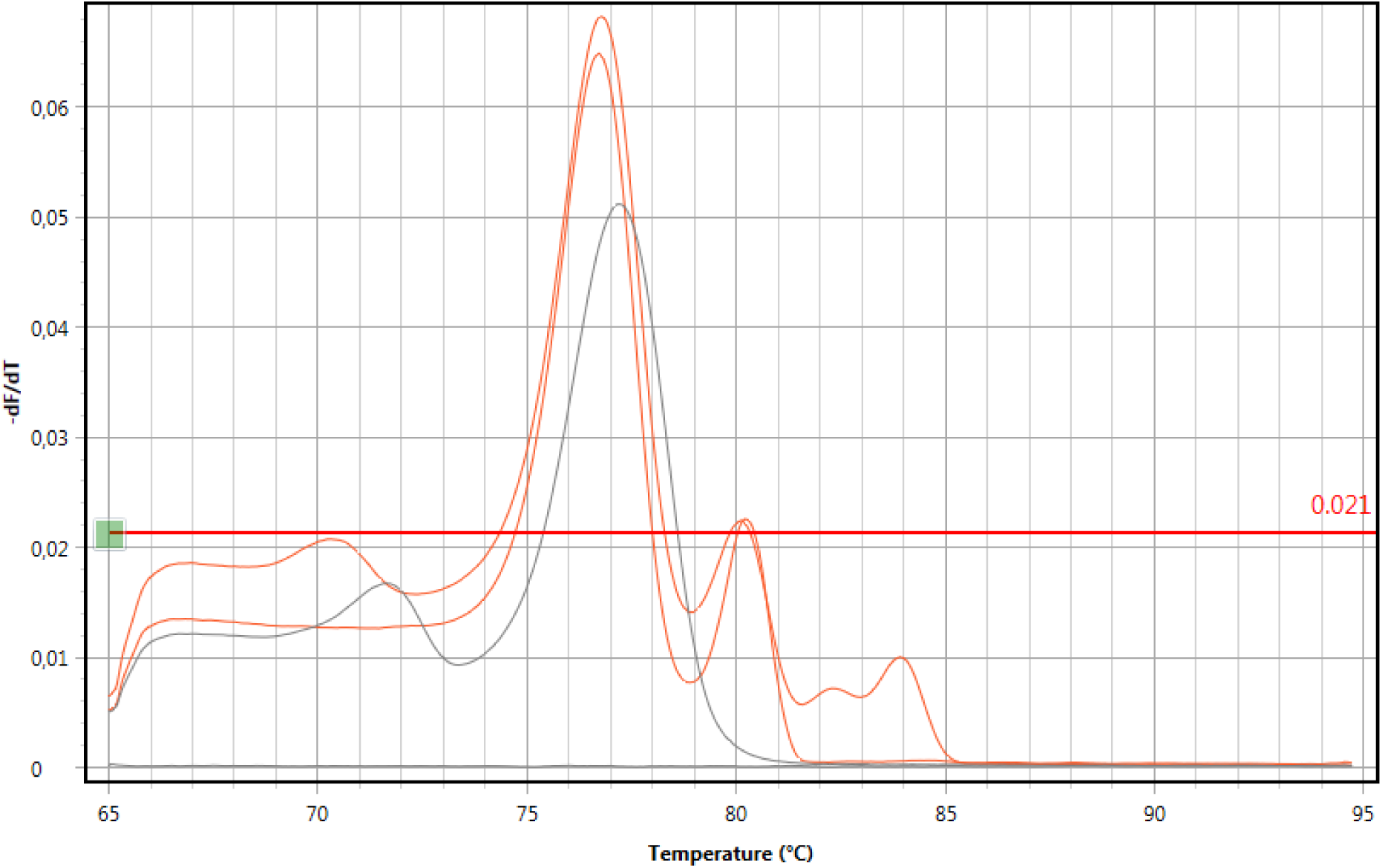
Melt curve of polymerase chain reaction of sample TLN18 (orange), no template control (grey) with preceding reverse transcription using primer pair Ec-2, each reaction in duplicate.

**Figure 6:**
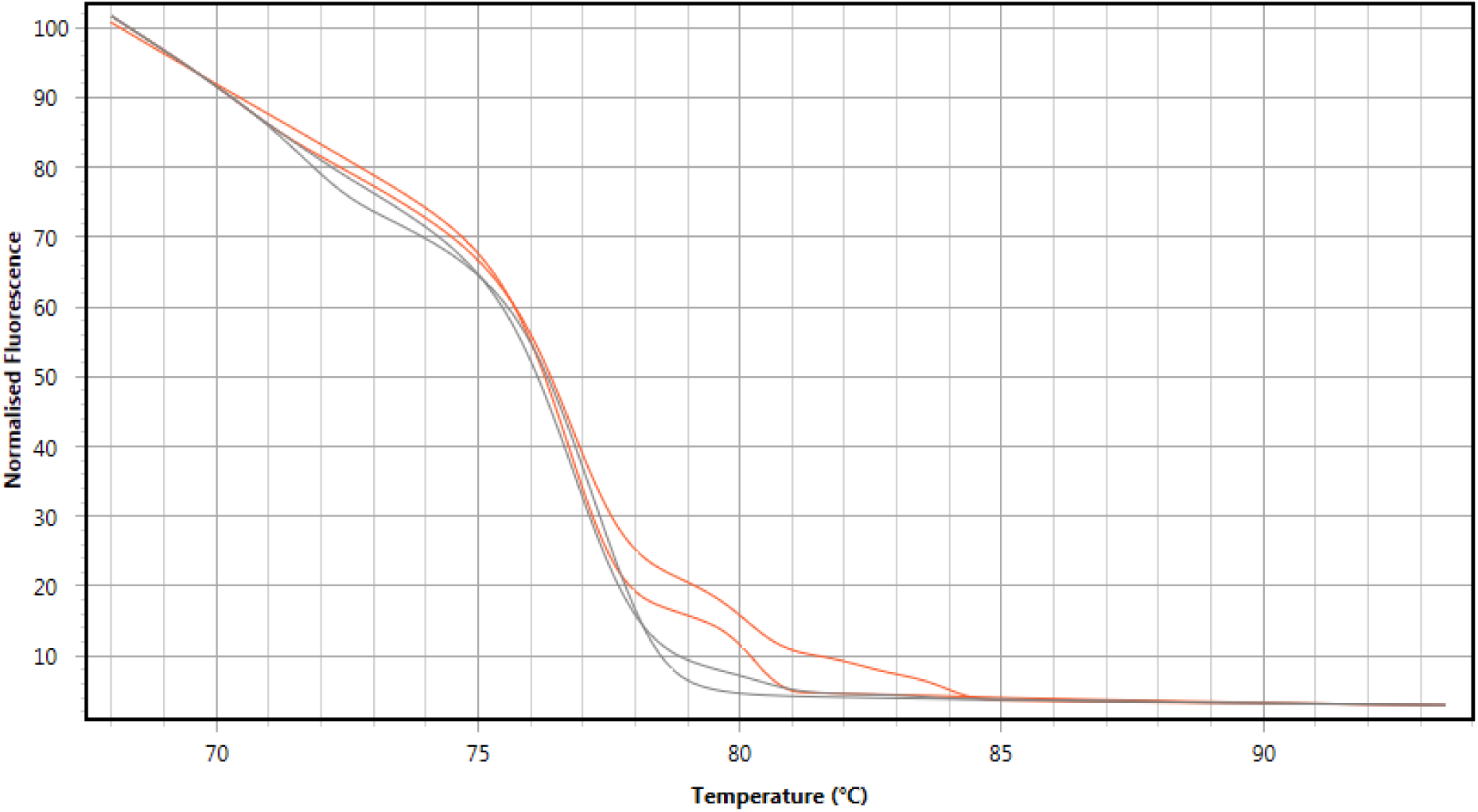
High resolution melt curve of polymerase chain reaction of sample TLN18 (orange), no template control (grey) with preceding reverse transcription using primer pair Ec- 2, each reaction in duplicate.

**Figure 7:**
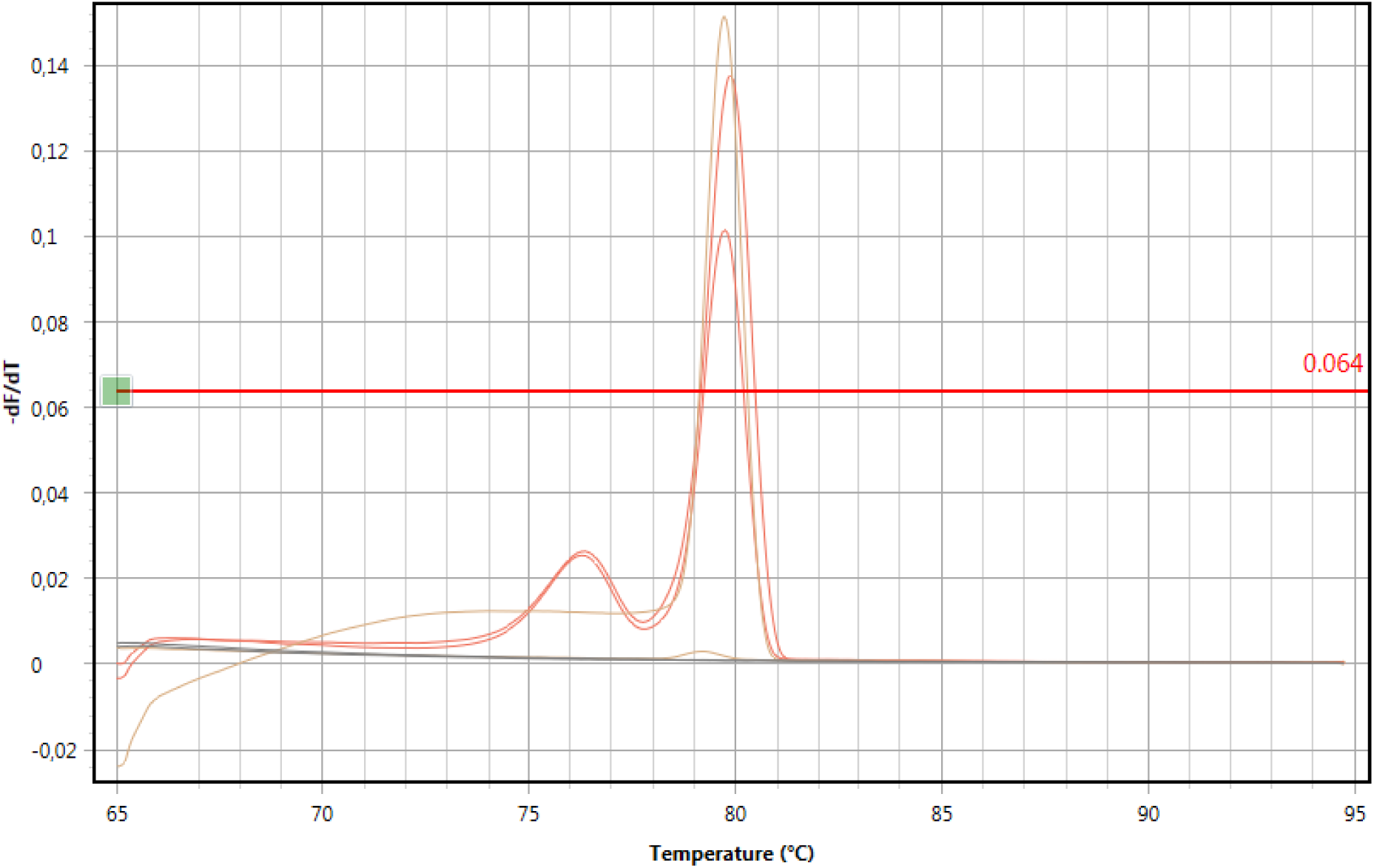
Melt curve of polymerase chain reaction of sample GL2 (orange), TLN18 (light brown), no template control (grey) without preceding reverse transcription using primer pair Ec-1, each reaction in duplicate.

**Figure 8:**
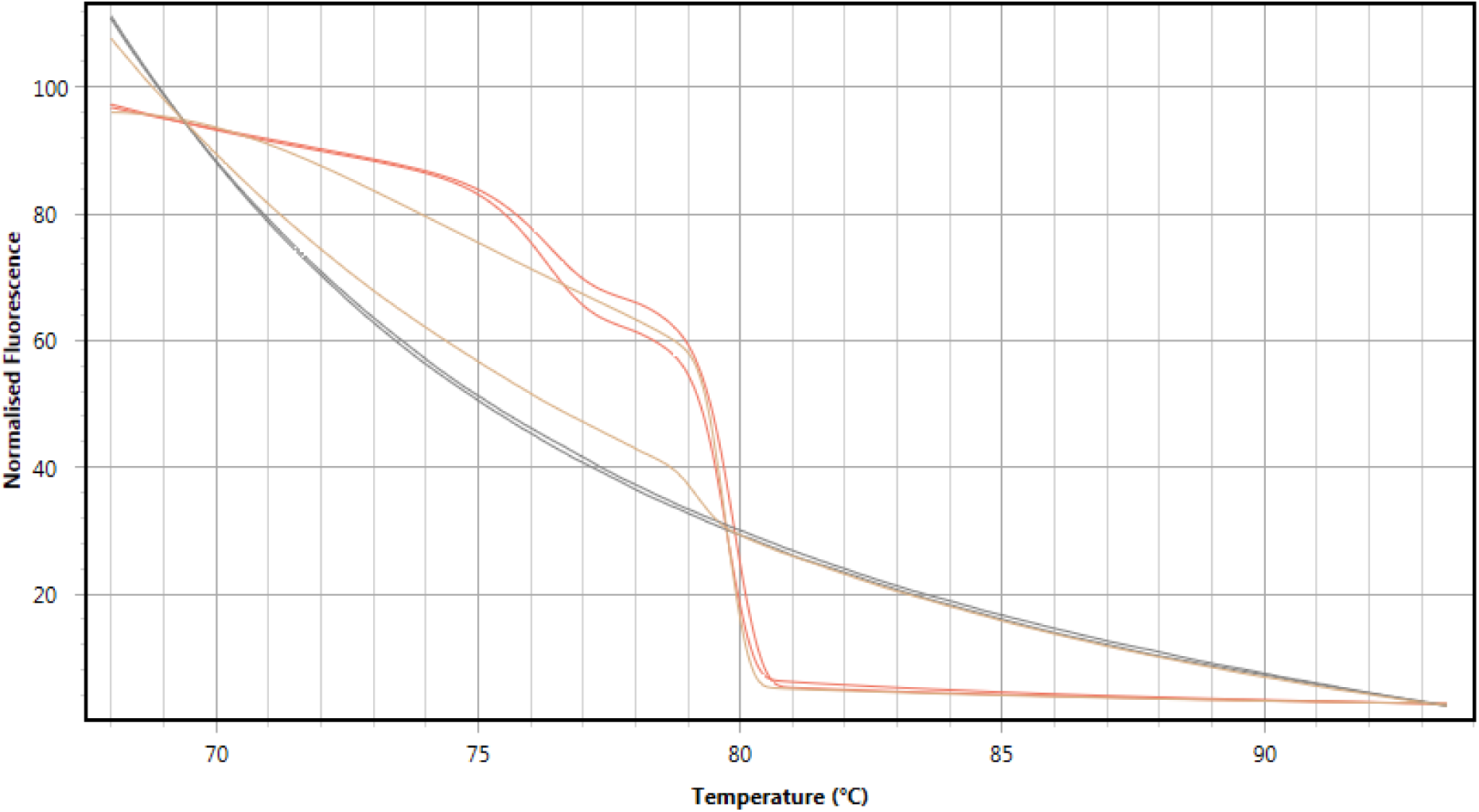
High resolution melt curve of polymerase chain reaction of sample GL2 (orange), TLN18 (light brown), no template control (grey) without preceding reverse transcription using primer pair Ec-1, each reaction in duplicate.

### 2.3 Samples from Kenya tested positive in real-time PCR assays and were confirmed by Sanger sequencing of the PCR products

Real-time PCR of two human hydatid cyst samples (M5 and M6 metacestode material of two different patients) and two fecal samples of carnivores (F4 dog, F3 unknown wild carnivore species) collected in Kenya and extracted with one DNA extraction method, resulted in detection of target genes of the genus *Echinococcus*. In both human metacestode samples (M5 and M6) the target region of NADH dehydrogenase subunit 5 (nad5, primer pair Ec-2) and cytochrome c oxidase subunit 1 (cox1, primer pair Ec-1) were detected. The carnivore feces samples (F3, F4-df, and F4-pFl) were determined as positive in the developed PCR assay targeting the Echinococcus gene nad5 (primer pair Ec-2). All respective PCR products showed specific amplification by melt curve and high resolution melt (HRM) curve analysis and were identified by Sanger sequencing as belonging to the species *E. granulosus* s.s. (samples M5, M6, and F3) and *E. canadensis* (samples F4-df and F4-pFl). Assay conditions (used primer pair, no previous reverse transcription), label for the resulting PCR product, melt curve analysis peak (*T*_*m*_), HRM curve analysis result, quantification cycle (*C*_*q*_), and the percentage of sequence identity of each amplicon with the respective sequence of *E. granulosus* s.s. or *E. canadensis* are listed in Table 7.

**Table 7:** Conditions and results of real-time PCR assays of samples from Kenya which tested positive and were confirmed by Sanger sequencing of the PCR products. Sample name (M5, M6, F3, F4-df, and F4-pFl, see details in Table 2 in Materials and Methods), primer pair (Ec- 1-FW/-REV or Ec-2-FW/-REV), sequence ID of PCR product, no reverse transcription (RT) prior to PCR performed (-), specific high resolution melting (HRM) curve profile determined (+), average temperature (°C) of melt curve analysis peak (*T*_*m*_), average quantification cycle *C*_*q*_), and percentage of sequence identity (% id seq) of PCR product with *E. granulosus* genotype G1 (E. gran. G1), G3 (E. gran. G3), or *E. canadensis* (E. can.).

| Sample | Primer set | Sequence ID | RT | HRM | $T_m$ (°C) | $C_q$ | % id seq<br>E. gran.<br>G1 | % id seq<br>E. gran.<br>G3 |
| --- | --- | --- | --- | --- | --- | --- | --- | --- |
| M5 | Ec-2 | B22 | - | + | 80.01 | 18.60 | 98.71 | 98.38 |
|  | Ec-1 | B20 | - | + | 79.78 | 26.28 | 98.64 | 97.83 |
| M6 | Ec-2 | B16 | - | + | 79.87 | 13.21 | 98.71 | 98.38 |
|  | Ec-1 | B15 | - | + | 79.71 | 23.88 | 98.64 | 97.83 |
| F3 | Ec-2 | B17 | - | + | 80.32 | 32.78 | 98.38 | 98.06 |
|  |  |  |  |  |  |  | % id seq E. can. |  |
| F4-df | Ec-2 | B23 | - | + | 79.85 | 21.78 | 99.68 |  |
| F4-pFl | Ec-2 | B18 | - | + | 80.35 | 35.62 | 99.68 |  |

One fecal sample from a domestic dog (F4) was processed by two methods. DNA was extracted directly from feces without any enrichment of taeniid eggs by using a commercial spin column kit (see method "ZM" described in detail in chapter Materials and Methods). This DNA template of fecal sample F4 was labelled F4-df. The other method, resulting in DNA template F4-pFl, attempted enrichment of taeniid eggs and purification from fecal material by employing an egg flotation method before extracting DNA from the eggs-in-water suspension also with the method "ZM". As indicated by a higher *C*_*q*_ value of 35.62, compared to a *C*_*q*_ value of only 21.78 for template F4-df, the amount of target DNA in template F4-pFl was quite low. This was reflected in the lack of a specific melt curve peak *T*_*m*_ and a specific HRM curve profile for one out of three identical PCR reaction set-ups using template F4-pFl (Figure 11 and Figure 12).

The respective graphs display the specific melt curve analysis peak *T*_*m*_ (9, 11, 13, and 15) and the specific HRM curve profile (10, 12, 14, and 16) in comparison with the *T*_*m*_ and HRM curve profile of the no-template controls (NTCs).

**Figure 9:**
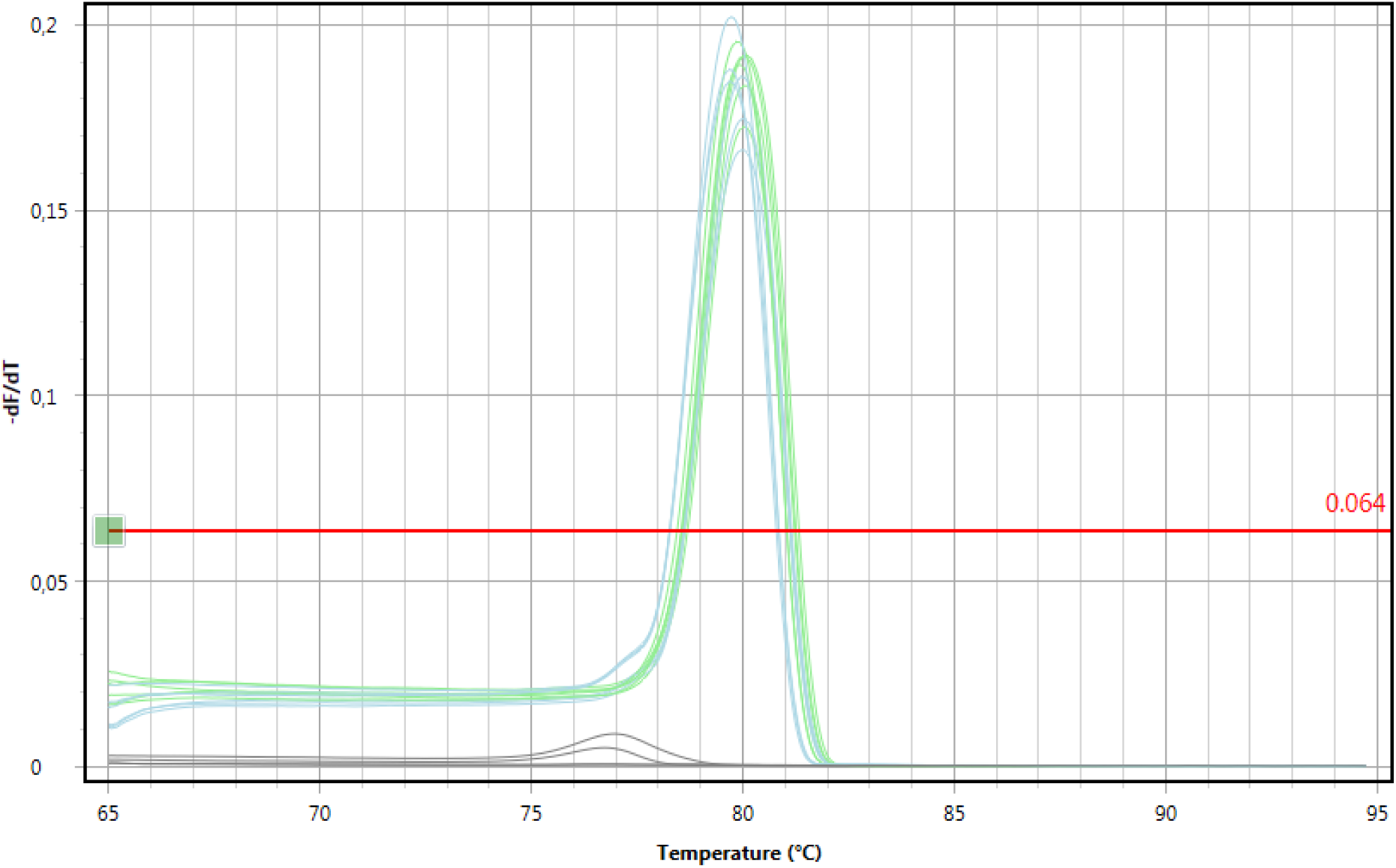
Melt curve of polymerase chain reaction of sample M5 (light green), F4-df (light blue), no template control (grey) without preceding reverse transcription using primer pair Ec- 2, each reaction in sextuplicate.

**Figure 10:**
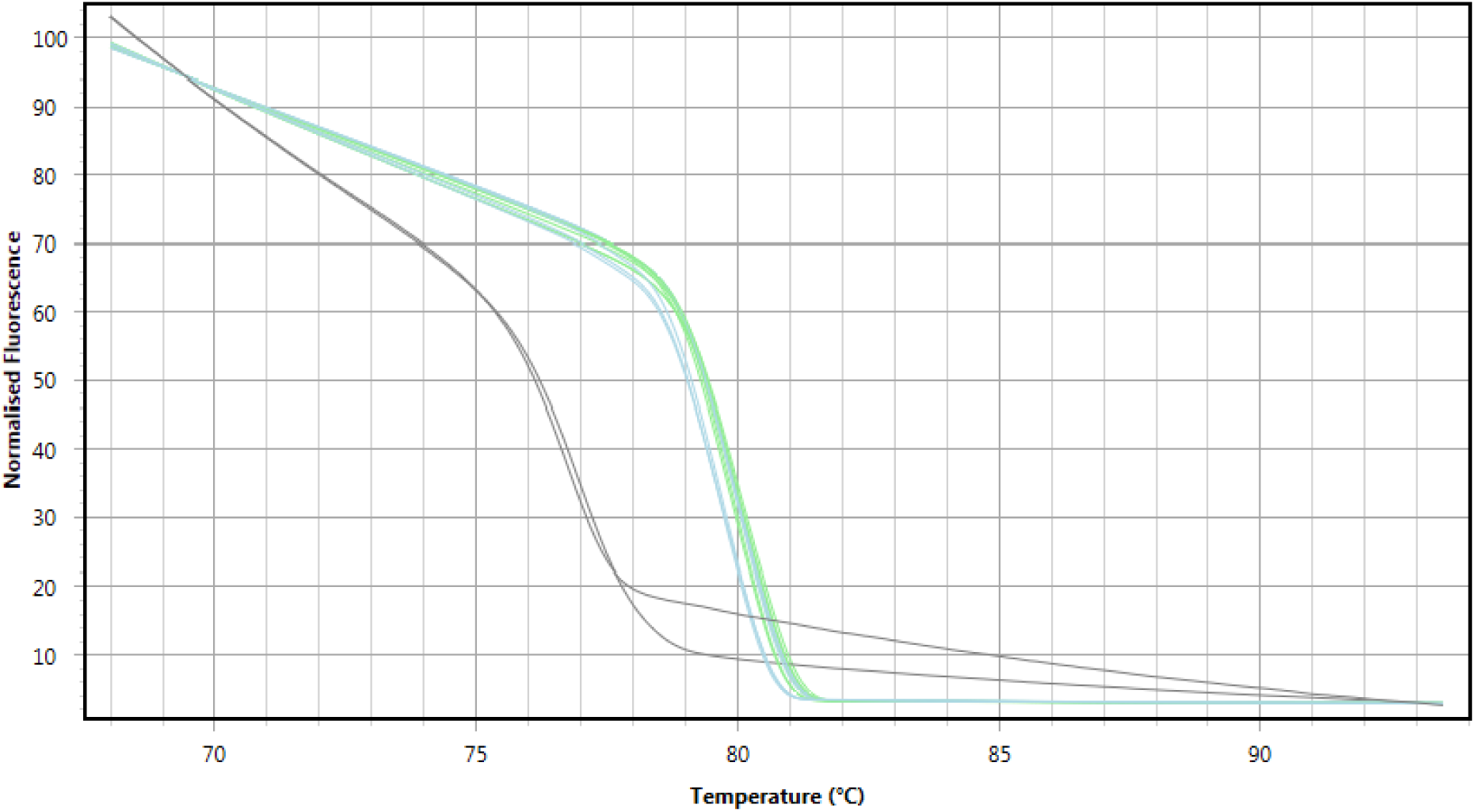
High resolution melt curve of polymerase chain reaction of sample M5 (light green), F4-df (light blue), no template control (grey) without preceding reverse transcription using primer pair Ec-2, each reaction in sextuplicate.

**Figure 11:**
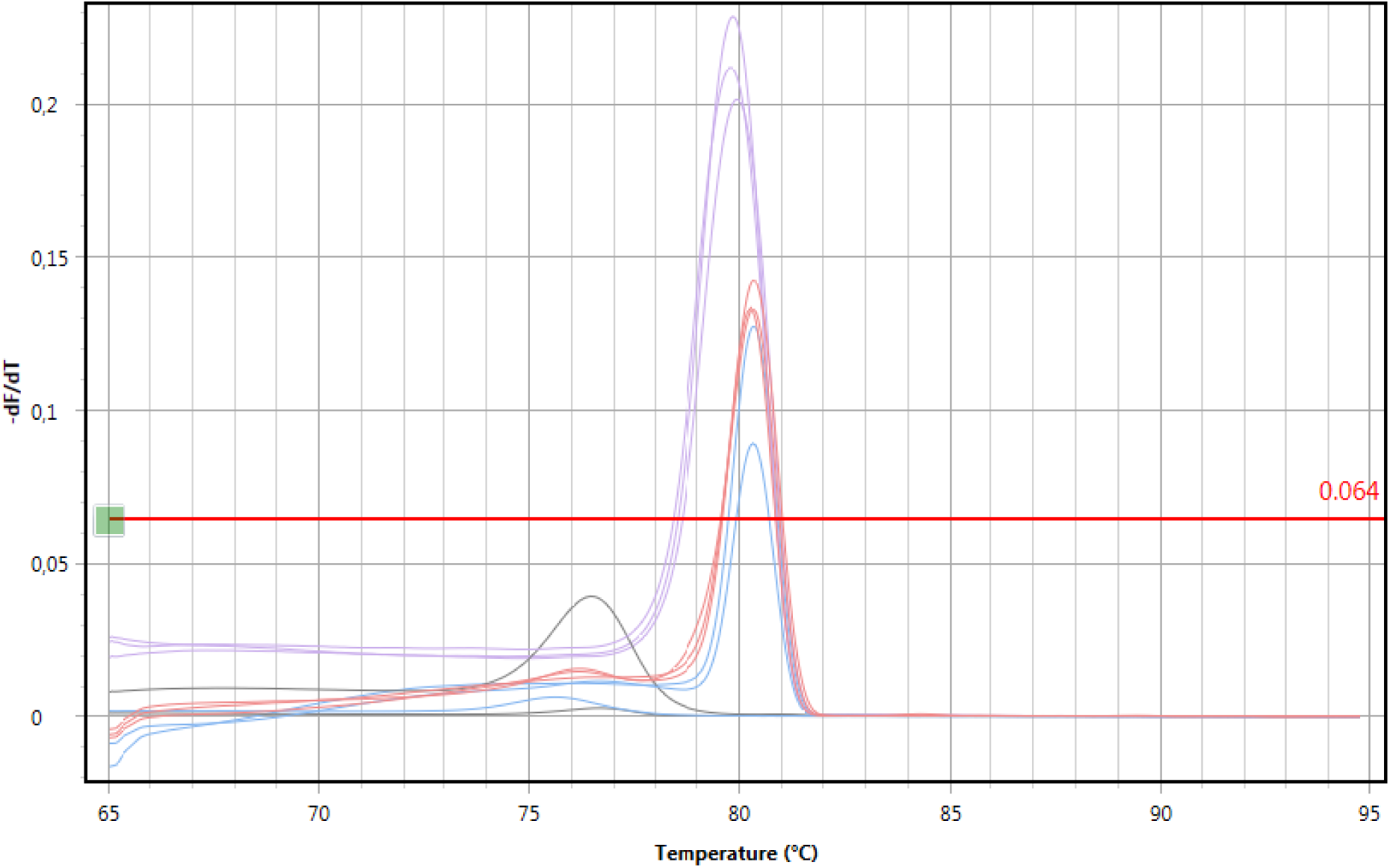
Melt curve of polymerase chain reaction of sample M6 (lavender), F3 (light red), F4-pFl (light blue), no template control (grey) without preceding reverse transcription using primer pair Ec-2, each reaction in triplicate.

**Figure 12:**
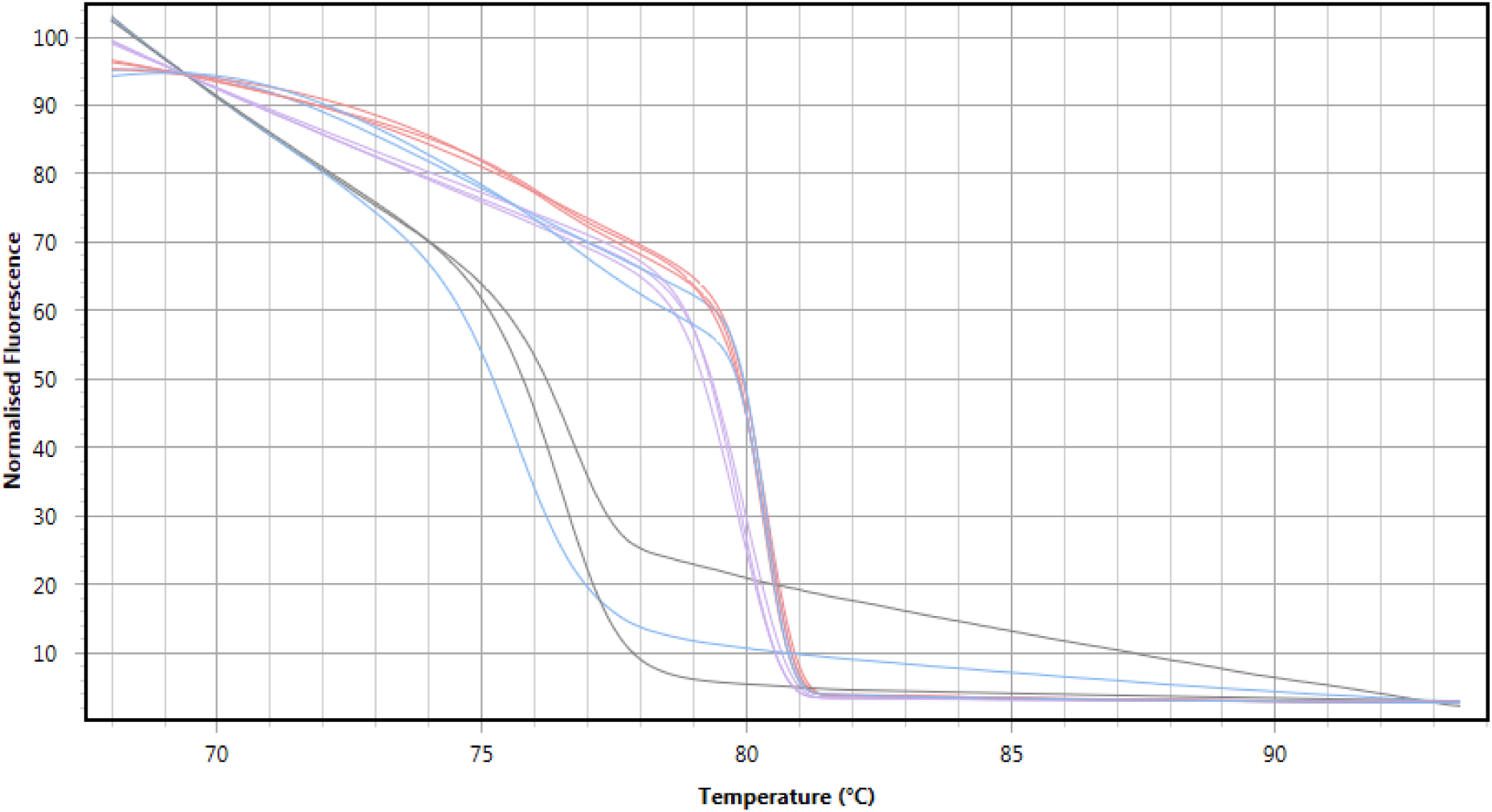
High resolution melt curve of polymerase chain reaction of sample M6 (lavender), F3 (light red), F4-pFl (light blue), no template control (grey) without preceding reverse transcription using primer pair Ec-2, each reaction in triplicate.

**Figure 13:**
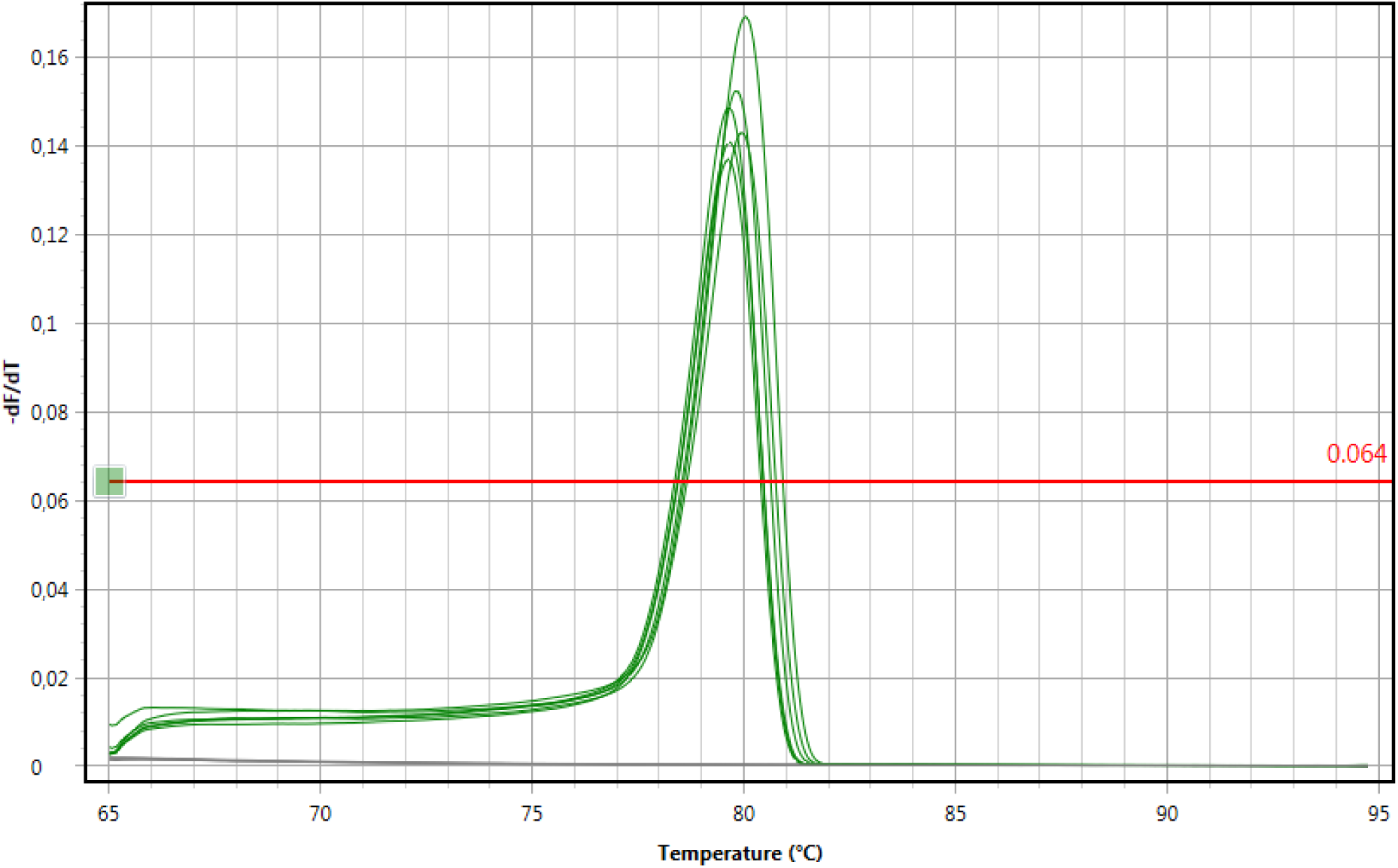
Melt curve of polymerase chain reaction of sample M5 (green), no template control (grey) without preceding reverse transcription using primer pair Ec-1, each reaction in sextuplicate.

**Figure 14:**
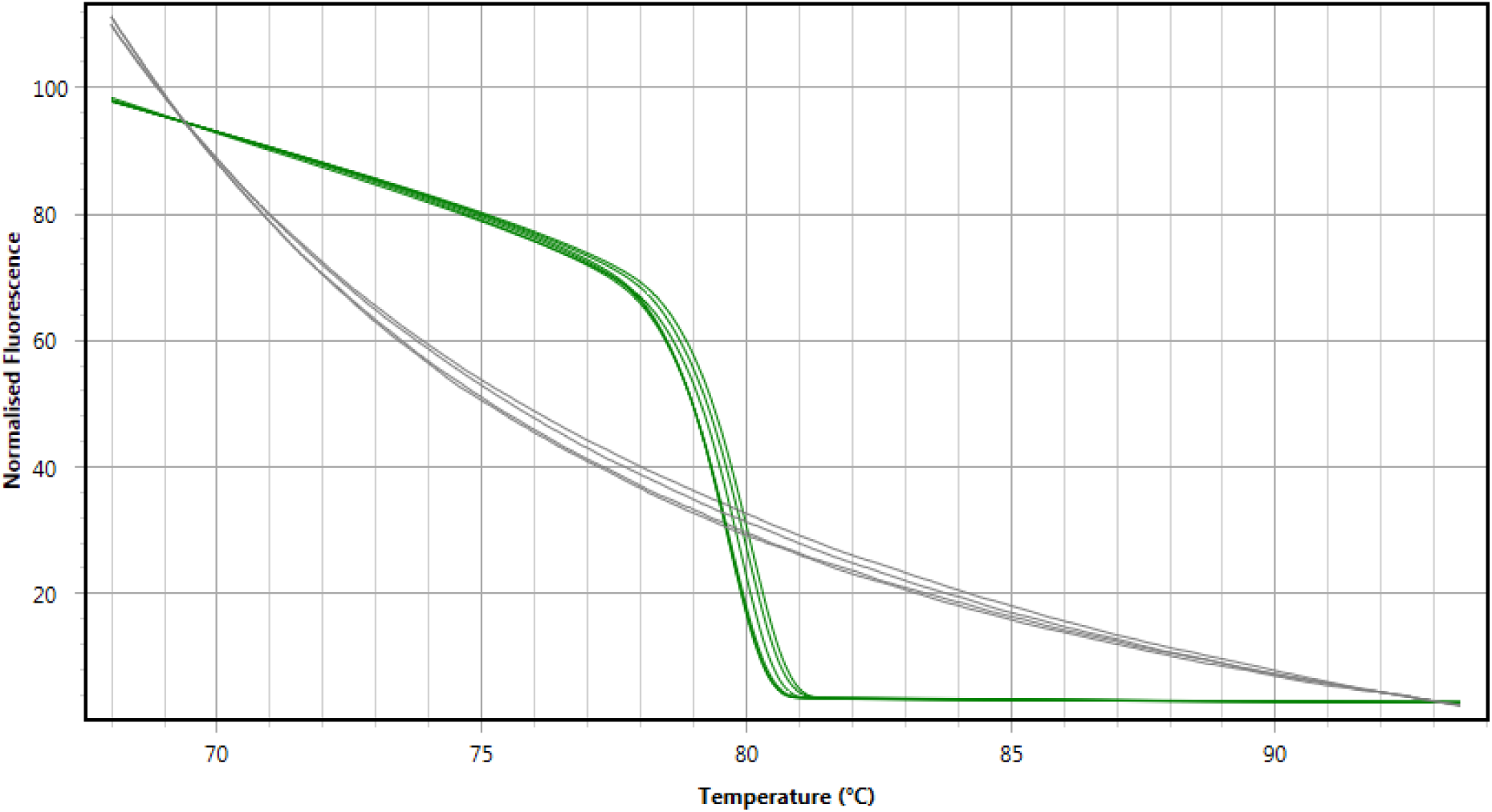
High resolution melt curve of polymerase chain reaction of sample M5 (green), no template control (grey) without preceding reverse transcription using primer pair Ec-1, each reaction in sextuplicate.

**Figure 15:**
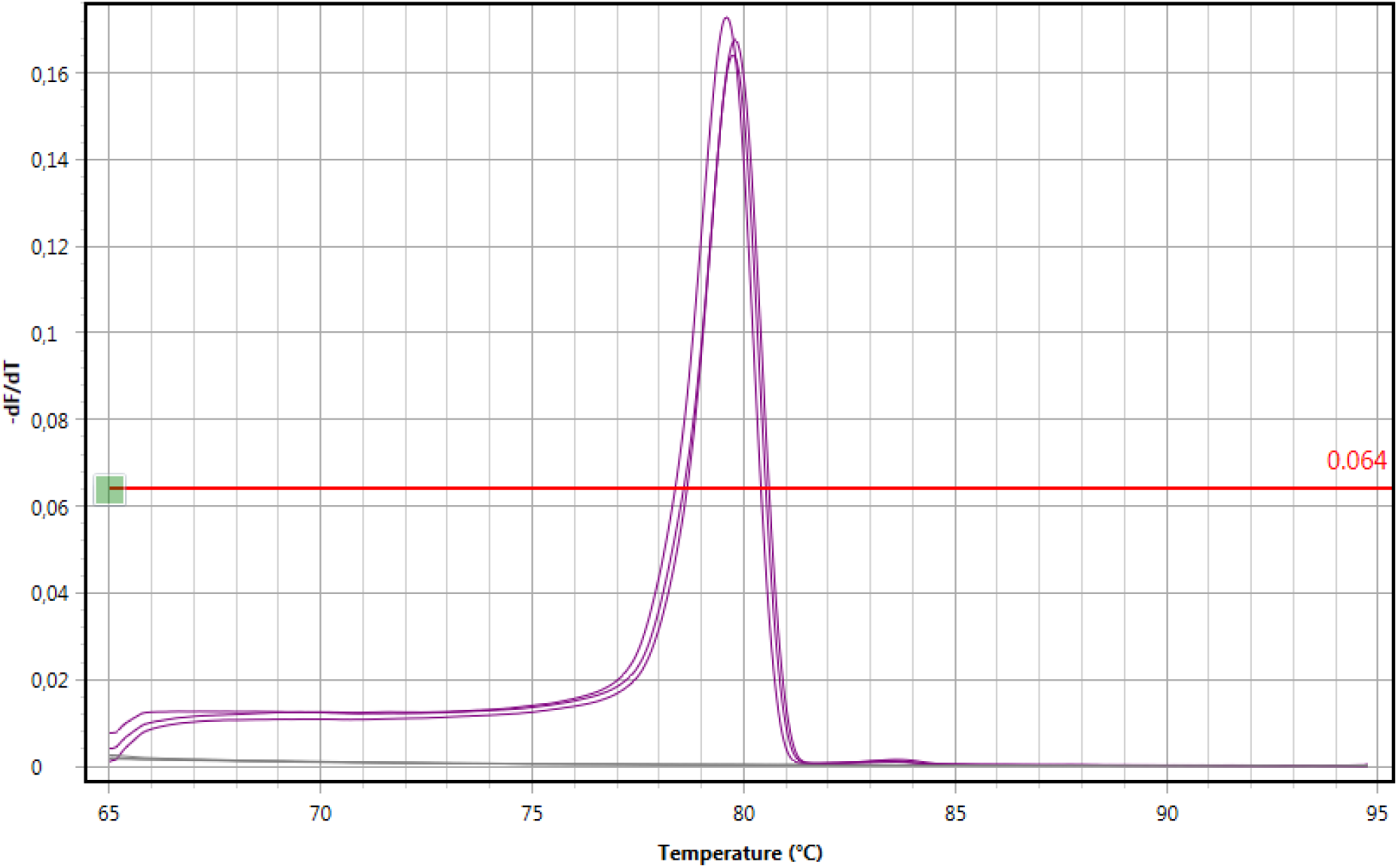
Melt curve of polymerase chain reaction of sample M6 (purple), no template control (grey) without preceding reverse transcription using primer pair Ec-1, each reaction in triplicate.

**Figure 16:**
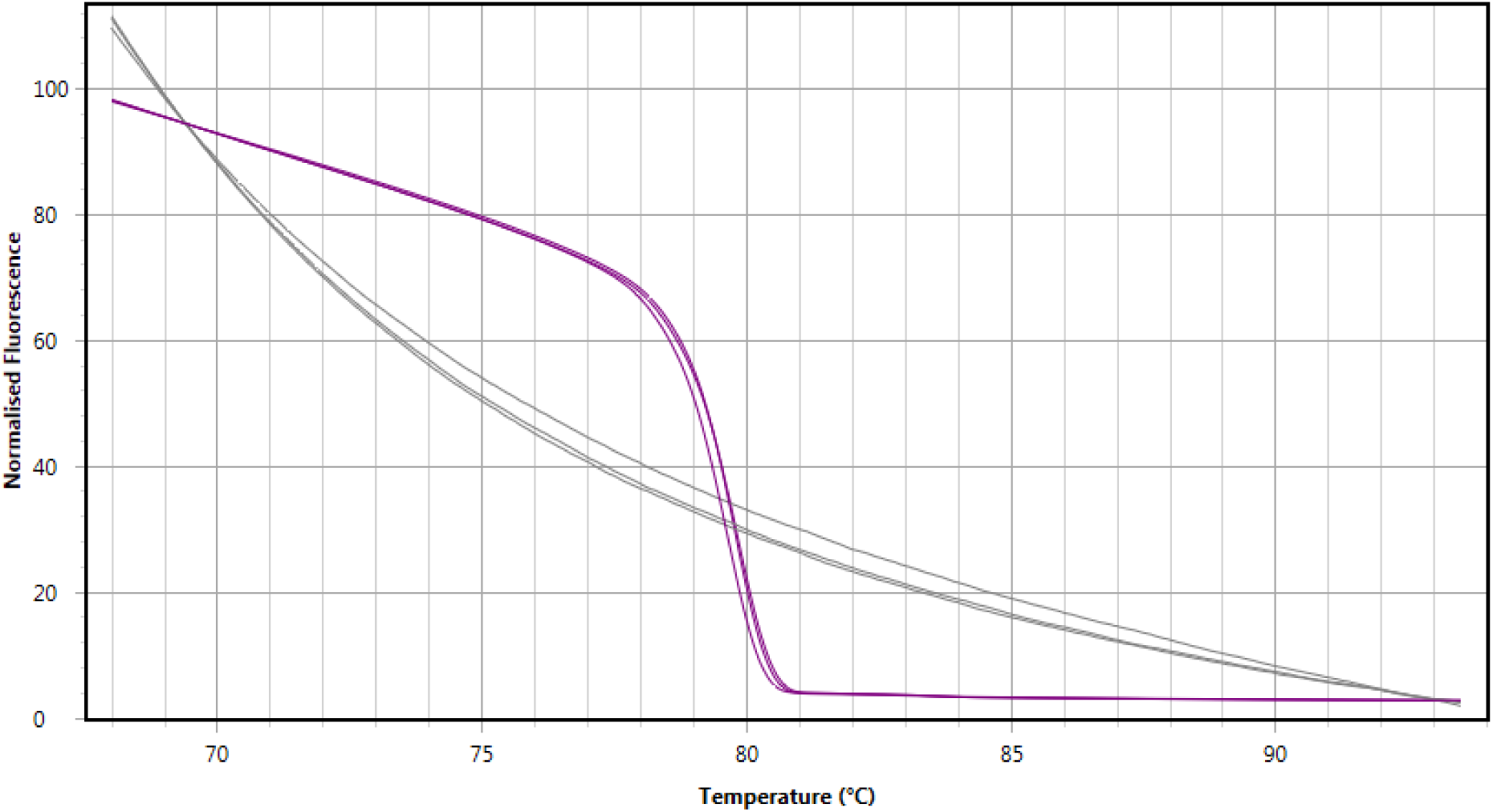
High resolution melt curve of polymerase chain reaction of sample M6 (purple), no template control (grey) without preceding reverse transcription using primer pair Ec-1, each reaction in triplicate.

In order to evaluate the replicability of the developed real-time PCR assay using primer pair Ec-2 and its potential to be extended from a qualitative to a quantitative molecular diagnostics method, six equal reactions were set up using DNA templates extracted from human hydatid cyst material (M5) and directly from a dog feces sample (F4-df). The graph of the corresponding cycling analysis of these PCR-assays is shown in Figure 17. For assays using template M5 an average quantification cycle (*C*_*q*_) value of 18.50 was determined. A standard deviation (χσn–1) value of only 0.33 of the *C*_*q*_ values for the six sample reactions indicates high replicability. In line with these results for the human hydatid cyst sample, also the standard deviation value of 0.13 which was determined for the *C*_*q*_ values of the six reactions using the direct DNA extract from a dog feces sample shows high replicability of the developed assay. The average *C*_*q*_ value of 21.89 for sample F4-df can still be considered as indicative for the presence of a quite high amount of target DNA compared to *C*_*q*_ values above 32 for other samples in this study, for which melt curve and HRM curve analysis have failed to show signs of specific amplification in some of the replicates.

Thus, these replicates of two reactions (for samples M5 and F4-df) can only prove high replicability of specific amplification and, therefore, the capability of reliable target detection of the developed PCR assay for medium to high ranges of target DNA concentration in samples. At this stage of the assay evaluation we cannot determine the sensitivity of the developed PCR assay. If a theoretical efficiency of 100% is assumed for specific target amplification, the average *C*_*q*_ values for samples M5 and F4-df indicate that the amount of target DNA in M5 is roughly 10 times higher than in F4-df. However, in order to quantify the unknown absolute target DNA concentration of a sample using a standard curve is essential.

**Figure 17:**
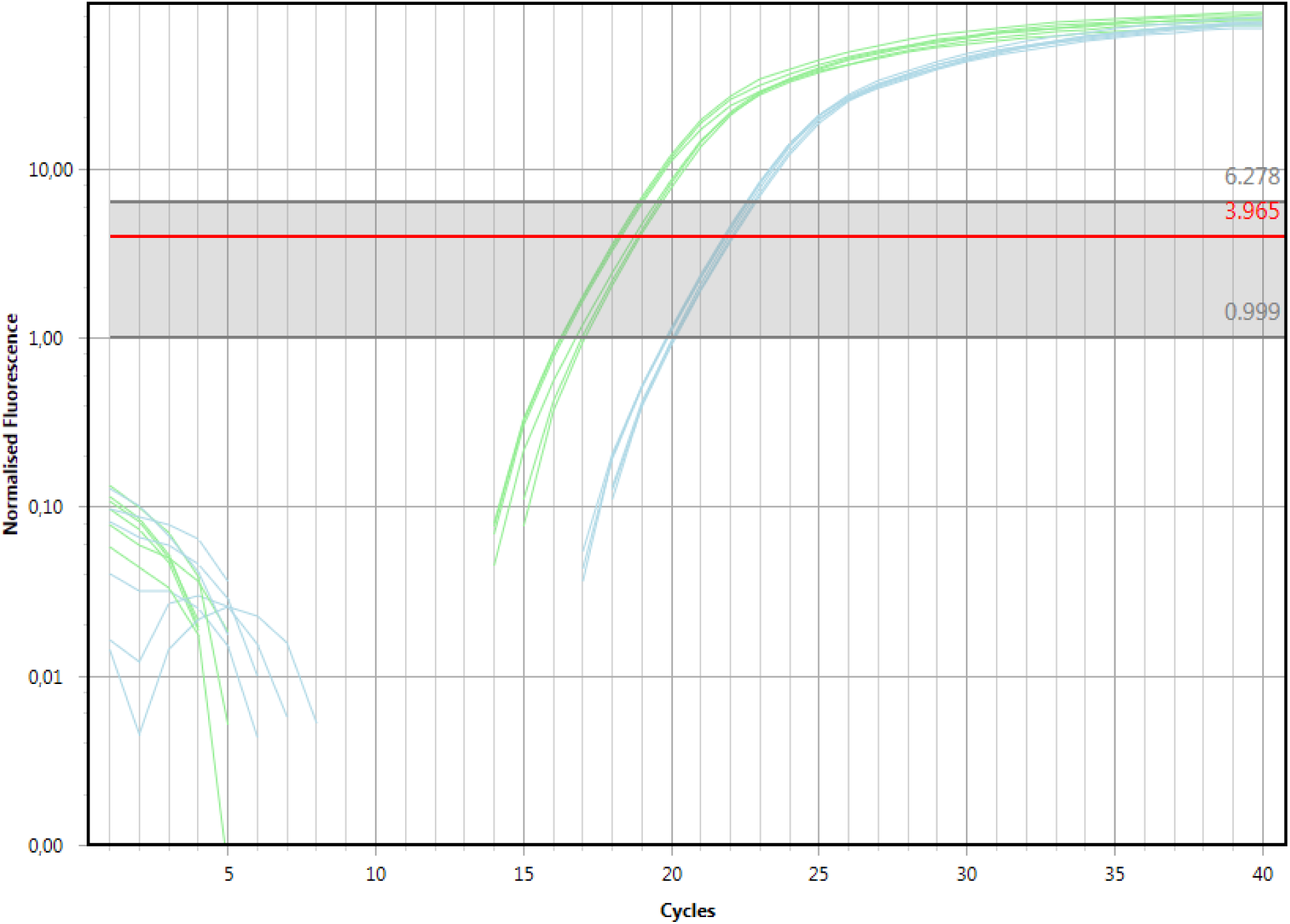
Cycling curve of polymerase chain reaction of sample M5 (light green), F4-df (light blue) without preceding reverse transcription using primer pair Ec-2, each reaction in sextuplicate.

### 2.4 Sequencing of PCR products of samples from Uganda confirmed molecular detection of *Echinococcus felidis*

The merged sequences B3 and B4 of PCR products of two lion feces samples from the Queen Elizabeth National Park in Western Uganda by primer pair Ec-1 were found to share the highest percentage of identity with the "lion strain" tapeworm species *E. felidis*. Sequence B3 (sample GL2) was completely identical with the predicted 372 bp long amplicon of primer pair Ec-1. This region of the full mitochondrial genome sequence of *E. felidis* (GenBank entry AB732958.1, genome position 10375 - 10746) contains parts of the gene cytochrome c oxidase subunit 1 (cox1) and t-RNA-Thr (trnT). With a value of 99.39% of identity also sequence B4 (sample TLN18) was almost identical with this region of the *E. felidis* mitochondrial genome.

The same Echinococcus species was also found to be the best match for all sequenced PCR products from Ugandan samples, when primer pair Ec-2 had been used for amplification. Sequence B1 (sample GL2, no reverse transcription before real-time PCR, RT-) was completely identical with the predicted 309 bp long amplicon of primer pair Ec-2. This region of the full mitochondrial genome sequence of *E. felidis* (GenBank entry AB732958.1, genome position 1054 - 1362) contains parts of the gene NADH dehydrogenase subunit 5 (nad5). When the target region of primer pair Ec-2 was amplified after reverse transcription (RT+) of the total DNA/RNA of sample GL2, still a high percentage of identity (98.71%) was found between the resulting sequence B8 and the respective region of the *E. felidis* mitochondrial genome. The same applied for the other fecal sample from a Ugandan lion (TLN18). Sequences B2 (RT-) and B5 (RT+) showed 99.35% and 96.76% of identity in pairwise alignments with the nad5 region of the mitochondrial genome of *E. felidis*, respectively.

The sequence B9 of the Ec-2 amplicon of a chest tissue sample (TLN5) of a giant forest hog (*Hylochoerus meinertzhageni*) from the Queen Elizabeth National Park in Western Uganda shared 97.68 % nucleotide bases with the corresponding nad5 region of *E. felidis*. Nucleic acid material of *E. felidis* has not been identified from tissue of this wild pig species before. While the sample had been labelled as "skin" material, it was reported that it had been excised from the chest region of a dead giant forest hog (personal communication, Siefert L., 2017). Therefore, this sample could have contained adjacent thoracic or abdominal organ tissue, such as of lungs or liver, harboring metacestode material. However, evidence of giant forest hogs as intermediate hosts for *E. felidis* rests on the identification of a single positive amplicon by the developed PCR assay, which calls for further data before drawing conclusions on the role of this wild pig species. The only other two mammalian species which had ever been identified as intermediate hosts of *E. felidis* by detection of nucleic acid sequences from metacestode material are one warthog in Uganda (Hüttner et al., 2009) and six hippopotami in South Africa (Halajian et al., 2017).

### 2.5 Sequencing of PCR products of samples from Kenya confirmed molecular detection of *Echinococcus granulosus* sensu stricto and *Echinococcus canadensis*

In contrast to all samples from Uganda, no sample from Kenya was found to contain nucleic acid sequences of *E. felidis*. Best matches of PCR product sequences for gene targets cox1 (primer pair Ec-1) and nad5 (primer pair Ec-2) for two human hydatid cyst samples (M5 and M6) and one wild carnivore fecal sample (F3) were found for *E. granulosus* s.s. genotype G1 (GenBank entry KY766891.1) and G3 (GenBank entry KY766892.1) with similar values ranging from 97.83% to 98,71% of identity.

PCR products B15 and B20 generated by primer pair Ec-1 of the two human hydatid cyst samples M6 and M5 shared a completely identical sequence. When primer pair Ec-2 had been used to amplify these templates the resulting amplicon sequences B16 and B22 still had 99.68% identity. Interestingly, these two sequences of human metacestode material were found to be very similar to the corresponding PCR product B17 of the carnivore fecal sample F3, with values of 99.68% and 99.35% for percentage of sequence identity with B16 and B22, respectively.

Independent of sample processing methods, the sequences of the Ec-2 amplicon of the fecal sample from a Kenyan dog (*Canis lupus familiaris*) (F4-df and F4-pFl) showed highest percentage of identity (99.68%) with the nad5 gene region of *E. canadensis* genotype G6 (GenBank entry AB208063.1). Interestingly, a high *C*_*q*_ value of 35.62 was determined, when DNA which had been extracted from taeniid eggs purified by a flotation method (see Materials and Methods) was used as template (F4-pFl). In comparison, when the same template volume was used in this real-time PCR assay but the DNA had been extracted directly from dog feces (F4-df), the observed *C*_*q*_ value was only 21.78. This indicates the presence of a higher amount of specific DNA target, when the direct method (F4-df) has been used. We would like to emphasize that the following comparison of DNA amount in different extracts from a sample can just be a rough estimate, because absolute quantification of target DNA amount of a sample by real-time PCR assays can only be achieved by previous evaluation of a standard curve. Assuming a theoretical efficiency of 100% of a PCR reaction, each cycle should double the amount of specific PCR product. A difference of 13.84 between the *C*_*q*_ values of the two extraction methods used for sample F4, therefore, can be estimated as roughly 10^4^ times more target DNA present in the template directly extracted from the fecal sample (F4-df) than in the template gained from extraction after egg flotation (F4-pFl). However, this significant difference in PCR product amount was not reflected in any sequence differences. Pairwise alignment of sequences B18 (F4-pFl) and B23 (F4-df) determined complete identity.

The region of the nad5 gene of *E. canadensis* genotype G6 (GenBank entry AB208063.1) which is targeted by the primer pair Ec-2 is identical with that of GenBank entry KC756275.1. While also assigned as genotype G6, this mitochondrial single gene sequence (nad5) from a sample from China was classified as belonging to the species *E. granulosus*. However, according to the latest consensus on taxonomy from phylogenetic studies (Addy et al., 2017), sequences closest to genotype G6 should rather be regarded as belonging to the species *E. canadensis*.

### 2.6 Data analysis of PCR product sequences of samples enabled distinct identification of *Echinococcus* species

All PCR products (for labels see Table 6 and 7) of primer pairs Ec-1 (B3, B4, B15, and B20) and Ec-2 (B1, B2, B5, B8, B9, B16, B17, B18, B22, and B23) which had indicated specific amplification in the developed real-time PCR assays were sequenced by using the Sanger sequencing method and underwent a consecutive BLAST search over the entire GenBank database. Overall, all carnivore fecal samples, animal and human hydatid cyst materials from Uganda and Kenya contained DNA sequences which matched best to a set of five GenBank entries representing three *Echinococcus* species *E. granulosus* s.s. (genotype G1 and G3, GenBank entries KY766891.1 and KY766892.1), *E. canadensis* (genotype G6, GenBank entry AB208063.1), and *E. felidis* (GenBank entry AB732958.1).

Formerly, *E. granulosus* and *E. canadensis* have been grouped under the taxonomic label *Echinococcus sensu lato* (s.l.), while today, genotypes G1-G3 of *E. granulosus* are considered to belong to the species *E. granulosus* sensu stricto (s.s.), and genotype G6/7 is regarded as a cluster of the species *E. canadensis* (Romig et al., 2015, Romig et al., 2017). The so-called "camel strain" *E. canadensis* genotype G6 (GenBank entry AB208063.1) shares an identical GenBank entry (GenBank entry KC756275.1) for the single gene NADH dehydrogenase subunit 5 (nad5), in which the nucleotide sequence is also assigned to the genotype G6, but is referred to as species *E. granulosus* instead of *E. canadensis*. Thus, we decided to identify any sequence matching to both entries as belonging to the species *E. canadensis*.

Pairwise alignment of the sequence of every single PCR product against each other amplicon and the best matching GenBank entry according to BLAST search resulted in two matrices of percentage of sequence identity (PID, in %) values, which are displayed in Table 8 for primer pair Ec-1 and Table 9 for primer pair Ec-2. When these PID values were visualized as plot diagrams for each primer pair, a clear distinction between lower and higher values was observed. For the PID values of primer pair Ec-1 (Figure 18) the lower PID value range was found to reach from 89.52 - 93.28% and the higher PID value range from 97.83 - 100%. The difference between the highest PID value of the low range and the lowest PID value of the high range was determined as 4.55%. The distinct separation of both PID ranges was visualized by a dotted line at half of that difference at a PID value of 95.56%.

Analogously, a clear distinction of PID values in two groups was found for sequences of PCR products generated by primer pair Ec-2 (Figure 19). Here, the lower PID value range was found to reach from 83.76 - 89.97% and the higher PID value range from 94.59 - 100%. The difference between the highest PID value of the low range and the lowest PID value of the high range was determined as 4.62%. The distinct separation of both PID ranges was visualized by a dotted line at half of that difference at a PID value of 92.28%.

For all of these PID values the following rule was found to apply. If the sequence of a PCR product matched best with the GenBank sequence for a certain *Echinococcus* species, the respective PID value fell into the higher range, while the PID values with sequences of all other *Echinococcus* species was only found in the lower range of PID values. Therefore, the developed PCR assays have shown the capability to unambiguously assign the sequence of a PCR product to one of the following *Echinococcus* species: *E. felidis*, *E. granulosus* s.s., or *E. canadensis*.

As no other species had been determined by sequencing the PCR products of the rather small sample set which had been examined here, it is not known, if the developed assays are able to detect and specifically amplify nucleic acid templates of further members of the genus *Echinococcus*. Furthermore, it awaits experimental evaluation using various field samples, if the herein observed clear species distinction would apply also for other *Echinococcus* species.

In order to estimate the potential sequence diversity of the two regions putatively amplified by primer pairs Ec-1 and Ec-2 of the developed PCR assays, we included non-redundant respective regions of several other *Echinococcus* species (see Table 10 for full list of corresponding GenBank entries). After multiple sequence alignment using Clustal Omega (http://ww.ebi.ac.uk/Tools/msa/clustalo/, Sievers et al., 2011), a phylogenetic tree was calculated based on average distance of PID using Jalview (version 2, last updated: 04 June 2014, Waterhouse et al., 2009). It should be noted that this type of basic tree cannot reflect a full phylogenetic analysis suitable for in-depth taxonomic or evolutionary conclusions of the genus *Echinococcus*. We rather intended to determine mere sequence diversity with respect to the potential of distinguishing possible PCR products of the developed real-time PCR assays by consecutive sequencing. Although the respective cox1 (Ec-1 primer pair, Figure 20) and nad5 (Ec-2 primer pair, Figure 21) mitochondrial genome regions of the *Echinococcus* species *E. equinus*, *E. ortleppi*, *E. shiquicus*, *E. oligarthrus*, and *E. vogeli* had been added to the multiple sequence alignments on which the phylogenetic tree calculations were conducted, the sequenced PCR products of the nucleic acid extracts from positive field samples clearly clustered only with *E. felidis*, *E. granulosus* s.s., and *E. canadensis*.

**Table 8:** Percentage of sequence identity (PID) values (%) of pairwise alignments of the sequence of every single PCR product of primer pair Ec-1 against each other amplicon and the best matching GenBank entry according to BLAST search.

| Gen-Bank ID/<br>Sequence label | AB732958 | B15 | B20 | B3 | B4 | KY766891 | KY766892 |
| --- | --- | --- | --- | --- | --- | --- | --- |
| AB208063 | 89,52 | 90,59 | 90,59 | 89,52 | 90,21 | 90,32 | 90,59 |
| AB732958 |  | 93,28 | 93,28 | 100,00 | 99,39 | 91,94 | 92,20 |
| B15 |  |  | 100,00 | 93,28 | 92,35 | 98,64 | 97,83 |
| B20 |  |  |  | 93,28 | 92,35 | 98,64 | 97,83 |
| B3 |  |  |  |  | 99,39 | 91,94 | 92,20 |
| B4 |  |  |  |  |  | 91,74 | 92,05 |
| KY766891 |  |  |  |  |  |  | 99,19 |

**Figure 18:**
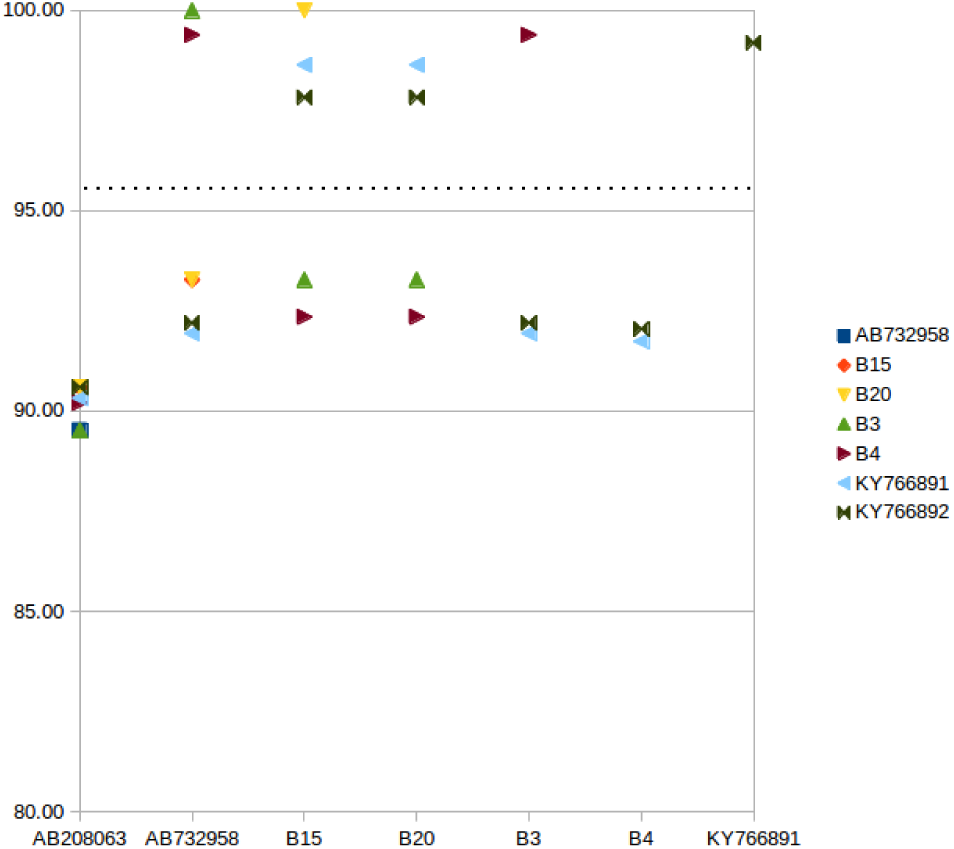
Plot of percentage of sequence identity (PID, in %) matrix for primer pair Ec-1 listed in Table 8.

**Table 9:** Percentage of sequence identity (PID) values (%) of pairwise alignments of the sequence of every single PCR product of primer pair Ec-2 against each other amplicon and the best matching GenBank entry according to BLAST search.

| GenBank ID/<br>Sequence label | AB732958 | B1 | B16 | B17 | B18 | B2 | B22 | B23 | B5 | B8 | B9 | KC756275 | KY766891 | KY766892 |
| --- | --- | --- | --- | --- | --- | --- | --- | --- | --- | --- | --- | --- | --- | --- |
| AB208063 | 88,18 | 88,18 | 89,35 | 89,64 | 99,68 | 87,54 | 89,03 | 99,68 | 85,67 | 87,22 | 84,39 | 100,00 | 88,71 | 89,03 |
| AB732958 |  | 100,00 | 88,29 | 87,97 | 88,78 | 99,35 | 87,97 | 88,75 | 96,76 | 98,71 | 97,68 | 88,46 | 87,03 | 87,97 |
| B1 |  |  | 88,29 | 87,97 | 88,78 | 99,35 | 87,97 | 88,75 | 96,76 | 98,71 | 97,68 | 88,46 | 87,03 | 87,97 |
| B16 |  |  |  | 99,68 | 89,68 | 88,18 | 99,68 | 89,64 | 86,62 | 87,58 | 84,70 | 89,35 | 98,71 | 98,38 |
| B17 |  |  |  |  | 89,97 | 87,86 | 99,35 | 89,94 | 86,31 | 87,26 | 84,70 | 89,64 | 98,38 | 98,06 |
| B18 |  |  |  |  |  | 87,86 | 89,35 | 100,00 | 85,99 | 87,54 | 84,39 | 99,68 | 88,39 | 88,71 |
| B2 |  |  |  |  |  |  | 87,97 | 88,10 | 96,12 | 98,71 | 97,30 | 87,82 | 87,03 | 87,66 |
| B22 |  |  |  |  |  |  |  | 89,32 | 86,31 | 87,90 | 84,33 | 89,03 | 98,71 | 98,38 |
| B23 |  |  |  |  |  |  |  |  | 85,94 | 87,50 | 84,33 | 99,68 | 88,35 | 88,67 |
| B5 |  |  |  |  |  |  |  |  |  | 96,12 | 94,59 | 85,94 | 85,35 | 86,26 |
| B8 |  |  |  |  |  |  |  |  |  |  | 96,91 | 87,22 | 86,16 | 87,11 |
| B9 |  |  |  |  |  |  |  |  |  |  |  | 84,70 | 83,76 | 84,50 |
| KC756275 |  |  |  |  |  |  |  |  |  |  |  |  | 88,71 | 89,03 |
| KY766891 |  |  |  |  |  |  |  |  |  |  |  |  |  | 99,03 |

**Figure 19:**
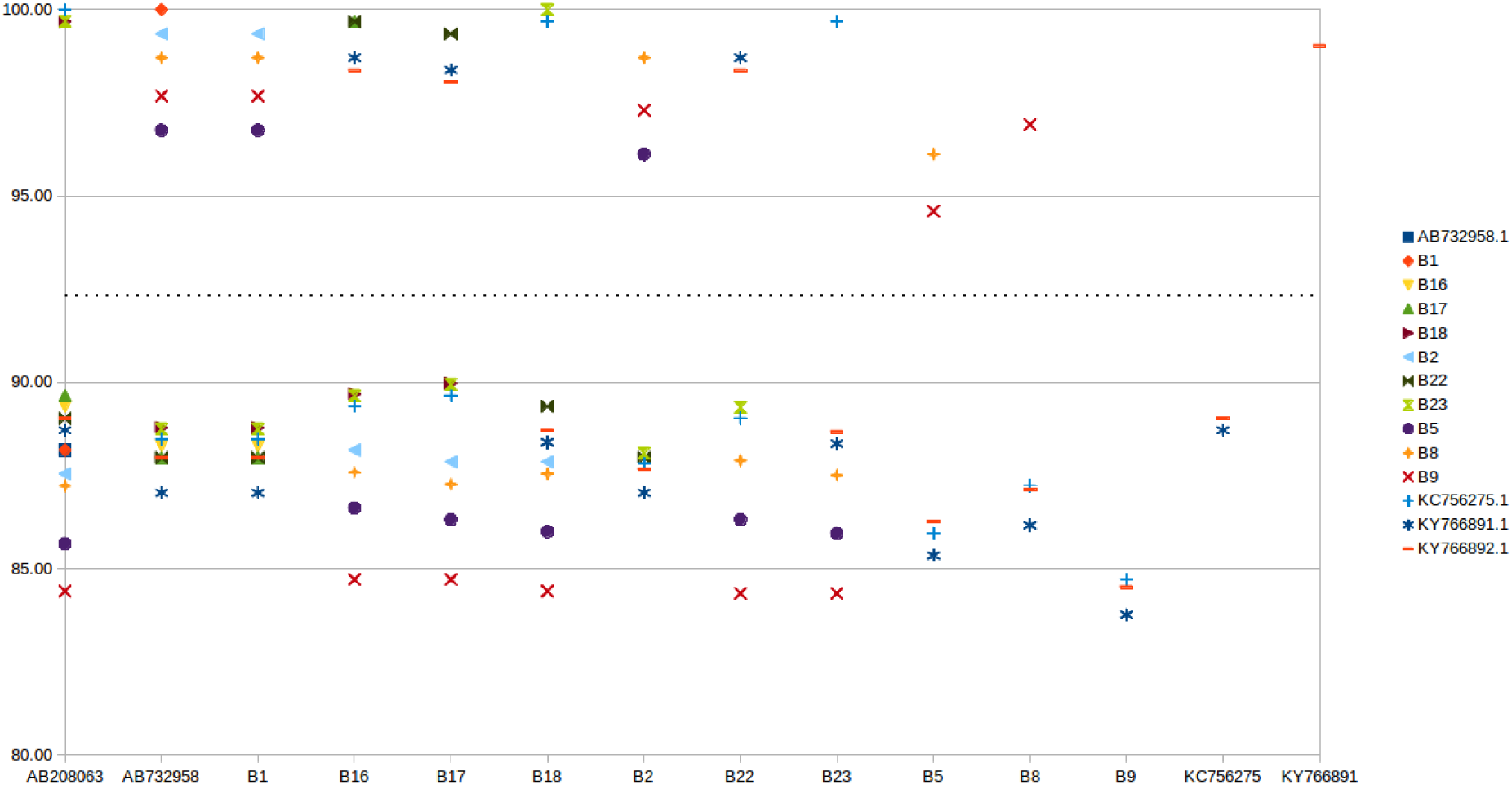
Plot of percentage of sequence identity (PID in %) matrix for primer pair Ec-2 listed in Table 9.

**Table 10:** GenBank entries of *Echinococcus* species used for constructing phylogenetic trees (Figure 20 and Figure 21), mitochondrial genome positions corresponding to the two regions targeted by primer pairs Ec-1 and Ec-2 in the sequence of *E. felidis*, isolate, host, GenBank taxon reference (Db xref taxon), developmental stage (Dev stage), country, and genotype or sample code, if given.

| GenBank ID | Ec-1 region | Ec-2 region | Organism | Isolate | Host | Db xref taxon | Dev stage | Country | Note |
| --- | --- | --- | --- | --- | --- | --- | --- | --- | --- |
| AB208063.1 | 10466-10837 | 1155-1463 | *E. canadensis* |  | camel | 519352 | protoscolex | Kazakhstan | genotype G6 |
| AB208064.1 | 10531-10904 | 1184-1498 | *E. shiquicus* |  | *Ochotona curzoniae* | 260967 | hydatid cyst | China |  |
| AB208545.1 | 10525-10903 | 1192-1503 | *E. oligarthrus* |  | *Mus musculus* | 6212 | protoscolex | Panama |  |
| AB208546.1 | 10498-10872 | 1173-1481 | *E. vogeli* |  |  | 6213 | hydatid cyst | Colombia |  |
| AB235846.1 | 10459-10830 | 1154-1462 | *E. ortleppi* |  | cattle | 446560 |  |  | genotype G5 |
| AB235847.1 | 10464-10835 | 1155-1463 | *E. canadensis* |  | pig | 519352 |  |  | genotype G7 |
| AB235848.1 | 10464-10834 | 1154-1462 | *E. canadensis* |  | moose | 519352 |  |  | genotype G8 |
| AB732958.1 | 10375-10746 | 1054-1362 | *E. felidis* |  | African lion | 460528 | egg | Uganda | sample code EfelUganda |
| AB745463.1 | 10465-10836 | 1154-1462 | *E. canadensis* | PA8695 | moose | 519352 | hydatid cyst | Finland | Genotype G10 |
| AB786664.1 | 10363-10731 | 1070-1378 | *E. granulosus* |  | *Homo sapiens* | 6210 |  | China Sichuan | sample code 52LI07 |
| AB786665.1 | 10355-10726 | 1049-1357 | *E. equinus* |  | *Equus caballus* | 519353 | hydatid cyst | United Kingdom | sample code EEQU/JEN |
| KC756275.1 | n.a. | 602-910 | *E. granulosus* | 40 | sheep | 6210 | metacestodes | China Sichuan | genotype G6 |
| KU601616.1 | 10334-10702 | 1046-1354 | *E. granulosus* |  | *Homo sapiens* | 6210 |  | Ethiopia | genotype Omo |
| KY766882.1 | 10249-10617 | 956-1264 | *E. granulosus* | ARG1 | cattle | 6210 |  | Argentina | genotype G1 |
| KY766891.1 | 10249-10617 | 956-1264 | *E. granulosus* | IND2 | buffalo | 6210 |  | India | genotype G1 |
| KY766892.1 | 10249-10617 | 956-1264 | *E. granulosus* | FRA2 | sheep | 6210 |  | France | genotype G3 |
| KY766893.1 | 10249-10617 | 956-1264 | *E. granulosus* | FRA1 | sheep | 6210 |  | France | genotype G3 |
| KY766897.1 | 10249-10617 | 956-1264 | *E. granulosus* | SPA4 | sheep | 6210 |  | Spain | genotype G3 |
| KY766904.1 | 10249-10617 | 956-1264 | *E. granulosus* | TUR2 | cattle | 6210 |  | Turkey | genotype G3 |

**Figure 20:**
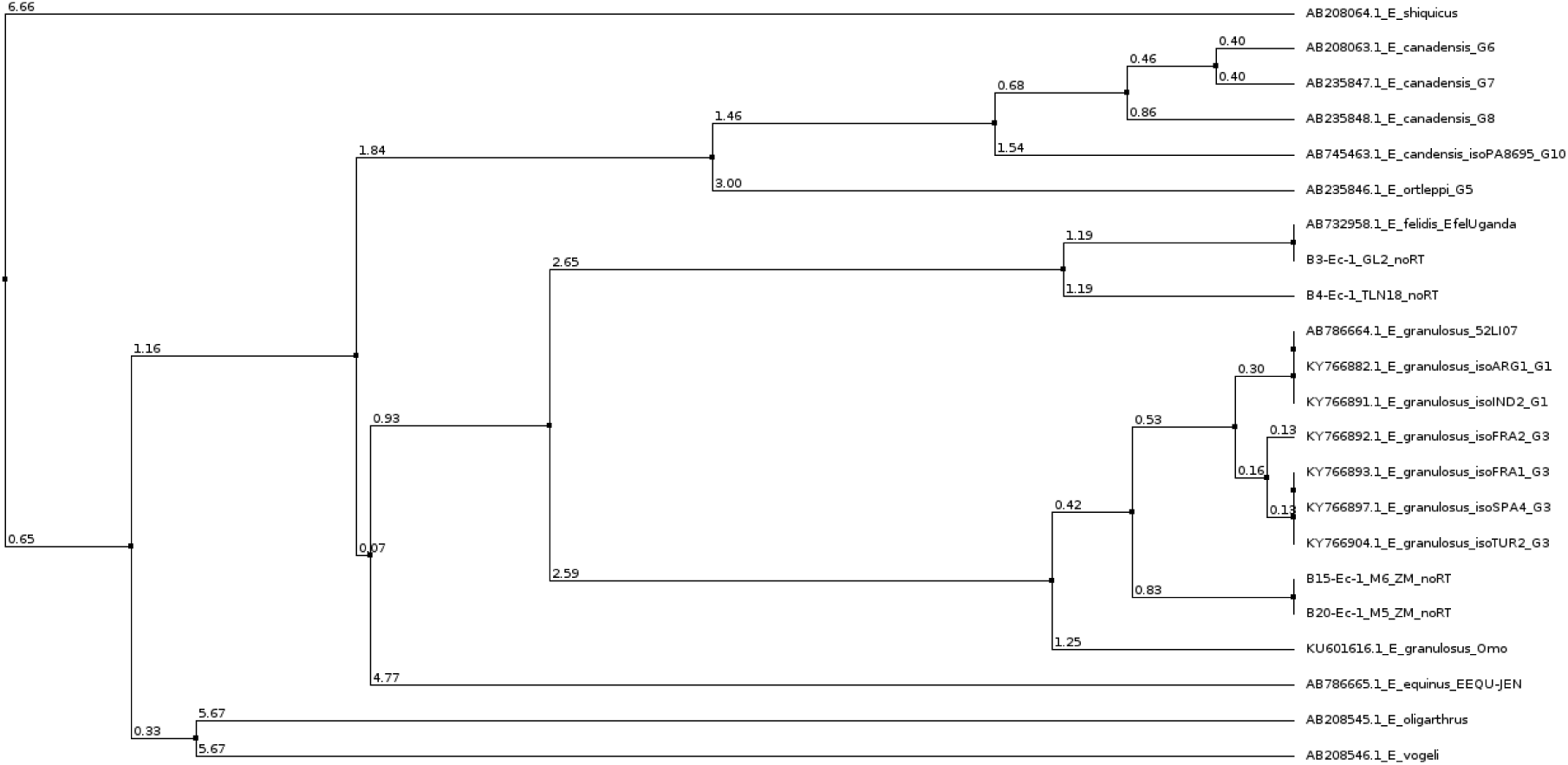
Phylogenetic tree of sequences of GenBank entries (Table 10) corresponding to the target region targeted by primer pair Ec-1 in the sequence of *E. felidis*; calculated based on average distance of percentage of identity (PID in %) between sequences using Jalview (version 2).

**Figure 21:**
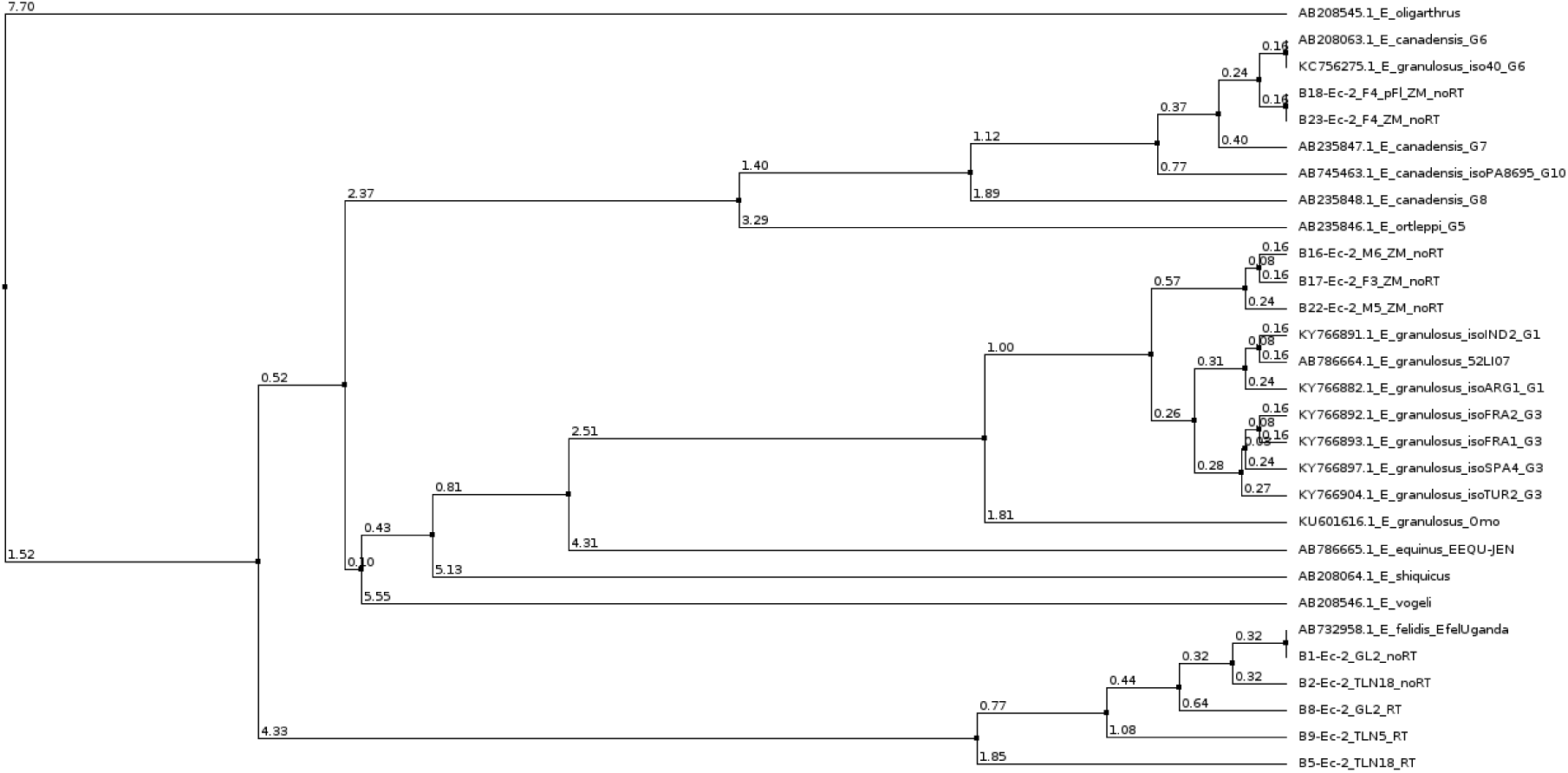
Phylogenetic tree of sequences of GenBank entries (Table 10) corresponding to the target region targeted by primer pair Ec-2 in the sequence of *E. felidis*; calculated based on average distance of percentage of identity (PID in %) between sequences using Jalview (version 2).

### 2.7 Preliminary application of developed PCR assays on blood samples did not detect nucleic acids of *Echinococcus* species

Apart from the samples, discussed in detail above, which were found positive with respect to signs of specific amplification in the developed real-time PCR assays and confirmed by consecutive sequencing, we performed a preliminary screening of several blood samples of various herbivore and carnivore wildlife species from Uganda. These samples included buffy coat preserved in 80% (vol/vol) ethanol and full blood with EDTA or Heparin. None of these blood samples was found positive for the presence of *Echinococcus* species. Some of these samples might have undergone several freeze-thawing cycles. Other of these samples had been stored for several years before processing. Therefore, some degree of nucleic acid degradation might apply for these samples. Furthermore, at the time these samples became available for processing, we did not have access to the TerraLyzer™ homogenization device.

Therefore, it could neither be shown nor ruled out that these real-time PCR assays might be capable of determining *Echinococcus* DNA in buffy coat or full blood samples from intermediate hosts (e.g. humans, herbivores) with cystic echinococcosis (CE) or definitive hosts (carnivores) infested with adult tapeworms. Examining this research question would require synchronized sampling of blood and hydatid cyst material from intermediate hosts or blood and feces from definitive hosts. Occasions for synchronized samplings of blood and microscopically confirmed parasite material, such as *Echinococcus* metacestodes from internal organ tissue or eggs from feces, could be abattoirs (livestock blood and tissue), necropsies of recently deceased wildlife (e.g. zebras, buffaloes, large carnivores blood and tissue), or routine vaccination programs (domestic dogs blood and feces). A less invasive approach for synchronized sampling of human CE patients (than surgery after diagnosis) could be ultrasonic mass screening of populations at risk and blood sampling of individuals with positive imaging results.

### 2.8 Differences in results of the developed PCR assays for samples from Uganda and Kenya require further evaluation with larger sample size

No systematic comparison between nucleic acid extraction methods could be performed, as only limited amounts of reagents and extraction kits were available for these preliminary studies on the proof of principle for real-time PCR detection of *Echinococcus* species gene targets from metacestode and egg nucleic acid material. However, for all samples from Kenya in which specific target amplification was determined by real-time PCR and was confirmed by Sanger sequencing of the PCR products, a uniform extraction method was applied (ZM, see chapter materials and methods for details). Two different methods (GL and TLN, see chapter materials and methods for details) were used for extraction of nucleic acid from two different fecal samples of two lions from Queen Elizabeth National Park in Uganda. Method TLN was also applied for processing tissue material from a giant forest hog from Queen Elizabeth National Park. As also all PCR products from Ugandan samples which indicated specific amplification were also confirmed by Sanger sequencing, extraction method bias probably does not explain the finding that the amplicons isolated from PCR reactions of all three Ugandan samples were uniformly determined as belonging to genome sequences of *E. felidis*, while three samples from Kenya were identified as containing material of *E. granulosus* s.s. and one sample material of *E. canadensis*. However, in fecal samples from lions in Kenya, E*. felidis* and *E. granulosus* s.s. had been found before (Kagendo et al., 2014. Geographical clustering or distribution patterns of *Echinococcus* species in definitive and intermediate host species could only be revealed by large scale screening of sufficient sample sizes. As the described nucleic acid extraction methods and real-time PCR assays have successfully passed this preliminary proof of principle study, they might be an ideal high-throughput method set for such an endeavor. However, it still has to be evaluated, if this molecular diagnostics protocol is less laborious, less hazardous, faster, and especially more sensitive and specific in comparison with other molecular or traditional methods, such as microscopic egg counting in feces or macro- and microscopic hydatid cyst inspection in intermediate host tissue.

### 2.9 Direct extraction of *Echinococcus* DNA from feces and consecutive one-step detection and quantification by real-time PCR assays could provide a robust, fast, and safe field method

As all of the samples which tested positive for the presence of *Echinococcus* species target genes had been preserved in 80% (vol/vol) ethanol at least several days at 4-8°C before nucleic acid extraction, this cheap and simple preservation method has proven to be suitable for consecutive diagnostics using real-time PCR assays. Some of these samples had even endured exposure to ambient tropical temperatures for three days during transport.

The intensity of infection of carnivores with adult *Echinococcus* tapeworms can be determined by egg counting in feces. However, processing of fecal samples for examination by light microscopy poses an occupational health risk to laboratory personnel. It had been shown for *Echinococcus multilocularis* that its eggs were killed by exposure to −83°C for 48 hours and to −196°C for 20 hours (Veit et al., 1995). But this type of deep-freezing devices or liquid nitrogen facilities is rarely available in local labs or remote regions of the study presented here.

While an initial lysis and homogenization step is also required for all nucleic acid extraction methods which we had used in fecal samples prior to performing real-time PCR assays, it reduces the exposure time to potentially infectious agents including other zoonotic pathogens, such as viruses and bacteria, considerably.

The monophasic solution of phenol and guanidine isothiocyanate, e.g. TRIzol ™ Reagent (Cat.no. 15596026, Thermo Fisher Scientific, Waltham, MA, USA) or TRI Reagent™ (R2050-1-200, Zymo Research Corp., Irvine, CA, USA), was found to reliably inactivate highly pathogenic avian influenza virus (HPAIV) and *Franciscella tularenis* bacterium (Hartman et al., 2012). Effective disinfection of mycobacteria (including *M. tuberculosis*) has long been understood to be especially difficult due to the waxy cell-walls of these bacteria. However, phenol, in a dilution of 5% only, was proven to be effective under all test conditions (Best et al., 1990).

The proprietary nucleic acid storage and extraction solution DNA/RNA Shield™ (R1100-250, Zymo Research Corp., Irvine, CA, USA) was proven to effectively inactivate viruses, bacteria, and yeast (Zymo Research Corp., http://www.zymoresearch.com/downloads/dl/file/id/502/r1100_50i.pdf, as accessed on 16 Sept 2017). After treatment of infectious or mock samples with DNA/RNA Shield™ for five minutes, titers of pathogens were determined by plaque assay or growth assay. The agents found to be inactivated by these experiments included *Influenza A virus*, *Zaire ebolavirus*, *Herpes simplex virus* 1 and 2, *Escherichia coli*, *Bacillus subtilis*, *Lactobacillus fermentum*, *Enterococcus feacalis*, *Listeria monocytogenes*, *Pseudomonas aeruginosa, Salmonella enterica*, *Staphylococcus aureus*, *Cryptococcus neoformans*, and *Saccharomyces cerevisiae*.

We found that different nucleic acid protocols worked independent of a specific lysis solution and that enrichment of eggs by flotation of fecal samples was not necessary. The efficiency of homogenization using locally sourced glass pearls and molecular-standard nuclease-free Zirconia beads was not compared directly by processing one sample using these two different lysis matrices. However, two samples assumed to be quite similar (both lion feces from QENP in 80% ethanol containing *E. felidis* eggs) were processed using local glass beads ("Gazaland mall", Kampala, Uganda) and Quick-RNA MiniPrep kit (R1054, Zymo Research Corp., Irvine, CA, USA) without DNase digest (sample GL2) or Zirconia beads and Directzol kit™ (Zymo Research Corp., Irvine, CA, USA) without DNase digest (sample TLN18). As listed in Table 6, real-time PCR runs of both samples using primer pairs Ec-1 and Ec-2 showed specific amplification according to melt curve and HRM curve analysis and displayed similar quantification cycle (*C*_*q*_) values.

DNA of several fecal and hydatid cyst samples (samples M5, M6, F3, F4-df, and F4-pFl) which tested positive for Echinococcus species were extracted by using ZymoBIOMICS™ lysis reagent (Zymo Research Corp., Irvine, CA, USA). According to the manufacturer's documentation, this reagent had been especially designed to process a wide range of tough-to-lyze samples including soil, feces, and tissue. While the ingredients of this proprietary solution by Zymo Research Corp. had not been published, it could be assumed that they denature proteins and solve lipid shells of pathogens, and therefore, inactivate them safely. Once tapeworm eggs had been lyzed and nucleic acids had been extracted, the templates to be used in PCR assays can be considered as low risk material to handle in the laboratory. Therefore, quantifying the infectious intensity of *Echinococcus* species in fecal samples by nucleic acid extraction and real-time PCR can be considered as less hazardous than determining the burden of *Echinococcus* eggs in carnivore feces by flotation methods followed by microscopic egg counting.

Quantitative real-time PCR (qPCR) allows not only to test for the presence of a given target sequence of a pathogen but also to determine the quantity of it in a sample (Wiedenmann et al., 1998, Le Cann, et al., 2004). Two examples of rough comparisons of the relative amount of *Echinococcus* target gene material in different samples are discussed in previous chapters 2.3. and 2.5. However, in order to allow the absolute quantification of a pathogen present in a sample, a real-time PCR assay requires an elaborate evaluation and validation procedure. Absolute quantification of a DNA target can be achieved directly by determining the number of replicates only in so-called digital PCR. However, real-time PCR, the method applied in the present study, requires establishment of a standard curve.

This standard curve has to be generated with samples of a known target DNA quantity over a wide range of concentrations. One commonly used method is to create plasmids containing the DNA target region, and produce a 10x dilution series over at least five steps. The exact DNA concentration in the starting dilution is determined by independent DNA quantification means, such as the spectrophotometric or fluorometric method (e.g. NanoDrop™ or Qubit™, Thermo Fisher Scientific, Waltham, MA, USA). From there, the known amount of DNA in each PCR reaction of the dilution series can be given as weight (e.g. ng) or converted to copy numbers of the target and is plotted along the x-axis against the measured *C*_*q*_ value along the y-axis. The unknown amount of target DNA in a sample can then be extrapolated by comparing the determined *C*_*q*_ value of the respective PCR reaction to the standard curve.

While the amount of DNA in a sample can be determined as weight or copy numbers using absolute quantification by real-time PCR, it is common practice, especially in the field of infectious disease research, to relate *C*_*q*_ values of samples to standard curves with biological units. For example, it was found for *Sudan ebolavirus* that the virus load measured in RNA copy numbers per ml of serum by RT-qPCR was consistently 3 - 4 log_10_ higher than the corresponding measurement by virus plaque forming units (PFU) per ml cell culture medium (Towner et al., 2004). This indicated that RNA copy numbers are only a measure of genomic molecules and not actual infectious virus particles. Analogously, the infectious unit regarding the potential zoonotic importance of *Echinococcus* species for human CE would be eggs in fecal samples of definitive hosts. Therefore, we suggest evaluating the correlation between target DNA copy numbers quantified by the developed real-time PCR assays in carnivore feces samples with the egg numbers counted by light microscopy after a fecal flotation procedure. As nested conventional end-point PCR has proven to be able to detect only one egg of *E. multilocularis* (Dinkel et al., 1998), we expect that the possible discrete resolution of a correlation between *C*_*q*_ value of the developed PCR assays and egg count lies at least within the range of few *Echinococcus* eggs.

While the first steps of the developed assays are highly practicable in field sampling and processing and do not require a sophisticated laboratory, up till recently, real-time PCR had been considered as difficult to perform under field conditions. Portable small real-time PCR thermocyclers, such as the Magnetic Induction Cycler micPCR™ system (Bio Molecular Systems, Potts Point, NSW, Australia), which had been used in our studies, offer some mobility. However, even this type of PCR instruments still requires constant supply of stable power at 230V. Recent attempts to miniaturize real-time PCR systems, in order to provide battery-powered portable devices by using microfluidics (Oblath et al., 2013, Ahrberg et al., 2016), could allow conducting all assay steps from nucleic acid extraction to result fast and directly in the field in near future.

### 2.10. Application of developed real-time PCR assays might have some advantages compared to other PCR methods with respect to sensitivity, prevention of false positives, and accurate intra- and interspecies identification of *Echinococcus* species

While there are several examples in literature which rank the sensitivity (in terms of limit of detection) of diagnostic PCR from highest to lowest in the order real-time PCR, nested PCR, single primer pair conventional/end-point PCR, this matter has been discussed controversially (Lemmon and Gardner, 2008, Bastien et al., 2008). The disparity of PCR methods in their respective sensitivity could often stem from different amplicon sizes. The shorter the length of the PCR product, the more efficiently and independent from any target degradation, it tends to be amplified. Real-time PCR products usually range from roughly 70 to 400 bp, while some conventional PCR assays had been designed to produce several hundreds to thousands of bp long amplicons. However, real-time PCR has some undisputed advantages over any conventional PCR method. Results can be literally displayed in real-time and are, therefore, faster than conventional PCR methods which require an agarose gel electrophoresis in order to visualize results. Sequencing of the PCR product is considered as gold standard for specific amplification. However, if combined with at least melt curve analysis and even better with high resolution melt curve analysis, real-time PCR is capable of a very accurate discrimination between specific or unspecific amplification.

In comparison, target DNA detection and species identification by using nested PCR and RFLP-fragment analysis requires two separate and consecutive PCR steps. For this purpose it is necessary to open the tubes of the first PCR reaction. If the target had been present in such a tube, the first PCR reaction resulted in high concentrations of the first specific amplicon.

This can lead to cross-contamination and false positive results in the second PCR reaction. RFLP-fragment analysis is cheaper than sequencing of the PCR product. While RFLP-fragment analysis has been evaluated for *E. granulosus* s.s., *E. felidis*, *E. equinus*, *E. ortleppi*, and *E. canadensis*, this method is not capable of identification of intraspecific variants and genotypes (Hüttner et al. 2008, 2009). On the contrary, sequencing of real-time as well as conventional one-step PCR products could be considered as more accurate in identifying the *Echinococcus* species and variant, if the inter- and intra-species diversity of the chosen amplicon is sufficiently high.

## 3. Conclusions and Outlook

We, hereby, present proof of principle for a fast (less or around two hours from sample to result), simple, and safe real-time PCR method for the detection of *Echinococcus* species in carnivore feces as well as in human patients and animal intermediate hosts affected by CE. Due to these features, this diagnostic method is predestined for future application in large-scale host population screenings and other high-throughput scenarios.

The developed real-time PCR assays can readily be extended from a mere qualitative to a quantitative method (real-time qPCR) by employing target DNA standard curve analysis. Determination of target copy numbers in fecal samples could help in predicting infectivity for intermediate hosts (including the zoonotic hazard potential for humans) and in monitoring deworming campaigns in definitive host populations. Likewise, quantification of copy numbers in hydatid cyst material of human CE patients (or of other intermediate hosts) could assist in the evaluation of therapy outcomes (surgical excision) and managing follow-up treatments (anthelmintic medication).

The developed real-time PCR assays were found to perform specific amplification of gene targets, when conducted as one-step RT-PCR, and, when run as PCR without preceding reverse transcription using total RNA and DNA extracts of samples as templates. However, no real-time RT-PCR has been contacted on DNase digested extracts containing total RNA only. Because of this, it has not been experimentally proven, if the developed assays are suitable for quantification of pure RNA molecules. In order to determine the infectivity and viability of *Echinococcus* species eggs and metacestodes by quantification of mRNA in fecal and hydatid cyst samples, further studies on expression patterns of genes in these different developmental stages of the parasites would be necessary. An example for this is a study on the expression profiles of oncospheres and early stage metacestodes of *E. multilocularis* (Huang et al. 2016). However, regarding *E. felidis*, RNA sequencing has not been conducted. While the complete mitochondrial genome of *E. felidis* has been deposited in GenBank (entry AB732958.1), only the whole nuclear genome sequences of the following two *Echinococcus* species are known to date: *E. multilocularis* (GenBank entry LN902841.1, Tsai et al., 2013) and *E. granulosus* (GenBank entry APAU02000001.1, Zheng et al., 2013). Knowledge of the complete nuclear genome and expression profiles of different developmental stages of *E. felidis* could inform targeted investigations on the infectivity and viability during this tapeworm's life cycle. This is especially important, as the complete range of definitive and intermediate hosts and the pathogenicity for humans of this tapeworm species are not known.

Real-time PCR has not been applied for the diagnosis of *E. felidis* before. The assays which we developed use real-time PCR in specifically detecting two distinct nucleotide sequence targets of the species *E. felidis* and two other *Echinococcu*s species. Apart from diagnosis of the "lion strain" tapeworm (*E. felidis*), detection of DNA material from *E. granulosus* s.s. and *E. canadensis* by the developed method was proven. The latter two species are endemic in East Africa and pose a considerable public health burden in this area.

In combination with Sanger sequencing the described real-time PCR assays showed their capability of clearly distinguishing between the species *E. felidis*, *E. granulosus* s.s, and *E. canadensis* in the samples from Uganda and Kenya which had tested positive for the presence of nucleic acid material of the genus *Echinococcus*. This is especially interesting, as this would allow examining the distinct full range of hosts and the specific potential of human pathogenicity for each of these three *Echinococcus* species, which are all endemic in the same geographical region, in large-scale screenings.

## References

Addy F, Alakonya A, Wamae N, Magambo J, Mbae C, Mulinge E, Zeyhle E, Wassermann M, Kern P, Romig T. Prevalence and diversity of cystic echinococcosis in livestock in Maasailand, Kenya. Parasitol Res. 2012 Dec;111(6):2289–94.

Addy F, Wassermann M, Kagendo D, Ebi D, Zeyhle E, Elmahdi IE, Umhang G, Casulli A, Harandi MF, Aschenborn O, Kern P, Mackenstedt U, Romig T. Genetic differentiation of the G6/7 cluster of Echinococcus canadensis based on mitochondrial marker genes. Int J Parasitol. 2017 Aug 3.

Ahrberg CD, Ilic BR, Manz A, Neužil P. Handheld real-time PCR device. Lab Chip. 2016 Feb 7;16(3):586–92.

Altschul SF, Gish W, Miller W, Myers EW, Lipman DJ. Basic local alignment search tool. J Mol Biol. 1990 Oct 5;215(3):403–10.

Alvarez Rojas CA, Romig T, Lightowlers MW. Echinococcus granulosus sensu lato genotypes infecting humans - review of current knowledge. Int J Parasitol. 2014 Jan;44(1):9–18.

Bastien P, Procop GW, Reischl U. Quantitative real-time PCR is not more sensitive than “conventional” PCR. J Clin Microbiol. 2008 Jun;46(6):1897–900.

Best M, Sattar SA, Springthorpe VS, Kennedy ME. Efficacies of selected disinfectants against Mycobacterium tuberculosis. Journal of Clinical Microbiology. 1990;28(10):2234–2239.

Bustin SA, Benes V, Garson JA, Hellemans J, Huggett J, Kubista M, Mueller R,Nolan T, Pfaffl MW, Shipley GL, Vandesompele J, Wittwer CT. The MIQE guidelines: minimum information for publication of quantitative real-time PCR experiments. Clin Chem. 2009 Apr;55(4):611–22.

Casulli A, Zeyhle E, Brunetti E, Pozio E, Meroni V, Genco F, Filice C. Molecular evidence of the camel strain (G6 genotype) of Echinococcus granulosus in humans from Turkana, Kenya. Trans R Soc Trop Med Hyg. 2010 Jan;104(1):29–32.

Dinkel A, von Nickisch-Rosenegk M, Bilger B, Merli M, Lucius R, Romig T. Detection of Echinococcus multilocularis in the definitive host: coprodiagnosis by PCR as an alternative to necropsy. J Clin Microbiol. 1998 Jul;36(7):1871–6.

Dinkel A, Njoroge EM, Zimmermann A, Wälz M, Zeyhle E, Elmahdi IE, Mackenstedt U, Romig T. A PCR system for detection of species and genotypes of the Echinococcus granulosus-complex, with reference to the epidemiological situation in eastern Africa. Int J Parasitol. 2004 Apr;34(5):645–53.

Halajian A, Luus-Powell WJ, Roux F, Nakao M, Sasaki M, Lavikainen A. Echinococcus felidis in hippopotamus, South Africa. Vet Parasitol. 2017 Aug 30;243:24–28.

Hartman A, Cole K, Homer L. Verification of Inactivation Methods for Removal of Biological Materials from a Biosafety Level 3 Select Agent Facility. Applied Biosafety. 2012;17(2).

Huang F, Dang Z, Suzuki Y, Horiuchi T, Yagi K, Kouguchi H, Irie T, Kim K, Oku Y. Analysis on Gene Expression Profile in Oncospheres and Early Stage Metacestodes from Echinococcus multilocularis. PLoS Negl Trop Dis. 2016 Apr 19;10(4):e0004634.

Hüttner M, Nakao M, Wassermann T, Siefert L, Boomker JD, Dinkel A, Sako Y, Mackenstedt U, Romig T, Ito A. Genetic characterization and phylogenetic position of Echinococcus felidis (Cestoda: Taeniidae) from the African lion. Int J Parasitol. 2008 Jun;38(7):861–8.

Hüttner M, Romig T. Echinococcus species in African wildlife. Parasitology. 2009 Sep;136(10):1089–95.

Hüttner M, Siefert L, Mackenstedt U, Romig T. A survey of Echinococcus species in wild carnivores and livestock in East Africa. Int J Parasitol. 2009 Sep;39(11):1269–76.

Kagendo D, Magambo J, Agola EL, Njenga SM, Zeyhle E, Mulinge E, Gitonga P, Mbae C, Muchiri E, Wassermann M, Kern P, Romig T. A survey for Echinococcus spp. of carnivores in six wildlife conservation areas in Kenya. Parasitol Int. 2014 Aug;63(4):604–11.

Koressaar T, Remm M. Enhancements and modifications of primer design program Primer3. Bioinformatics. 2007 May 15;23(10):1289–91.

Le Cann, P., S. Ranarijaona, S. Monpoeho, F. Le Guyader, and V. Ferre. 2004. Quantification of human astroviruses in sewage using real-time RT-PCR. Res. Microbiol. 155:11–15.

Lemmon GH, Gardner SN. Predicting the sensitivity and specificity of published real-time PCR assays. Ann Clin Microbiol Antimicrob. 2008 Sep 25;7:18.

Magambo J, Njoroge E, Zeyhle E. Epidemiology and control of echinococcosis in sub-Saharan Africa. Parasitol Int. 2006;55 Suppl:S193–5.

Mbaya H, Magambo J, Njenga S, Zeyhle E, Mbae C, Mulinge E, Wassermann M, Kern P, Romig T. Echinococcus spp. in central Kenya: a different story. Parasitol Res. 2014 Oct;113(10):3789–94.

Michener CD, Sokal RR. A quantitative approach to a problem of classification. Evolution 1957;11:490–499.

Morrison TB, Weis JJ, Wittwer CT. Quantification of low-copy transcripts by continuous SYBR Green I monitoring during amplification. Biotechniques. 1998 Jun;24(6):954–8, 960, 962.

Mutwiri T, Magambo J, Zeyhle E, Mkoji GM, Wamae CN, Mulinge E, Wassermann H, Kern P, Romig T. MOLECULAR CHARACTERISATION OF ECHINOCOCCUS GRANULOSUS SPECIES/STRAINS IN HUMAN INFECTIONS FROM TURKANA, KENYA. East Afr Med J. 2013 Jul;90(7):235–40.

Nakao M, Lavikainen A, Iwaki T, Haukisalmi V, Konyaev S, Oku Y, Okamoto M, Ito A. Molecular phylogeny of the genus Taenia (Cestoda: Taeniidae): proposals for the resurrection of Hydatigera Lamarck, 1816 and the creation of a new genus Versteria. Int J Parasitol. 2013 May;43(6):427–37.

Oblath EA, Henley WH, Alarie JP, Ramsey JM. A microfluidic chip integrating DNA extraction and real-time PCR for the detection of bacteria in saliva. Lab Chip. 2013 Apr 7;13(7):1325–32.

Omer RA, Dinkel A, Romig T, Mackenstedt U, Elnahas AA, Aradaib IE, Ahmed ME, Elmalik KH, Adam A. A molecular survey of cystic echinococcosis in Sudan. Vet Parasitol. 2010 May 11;169(3–4):340–6.

Romig T, Ebi D, Wassermann M. Taxonomy and molecular epidemiology of Echinococcus granulosus sensu lato. Vet Parasitol. 2015 Oct 30;213(3–4):76–84.

Romig T, Deplazes P, Jenkins D, Giraudoux P, Massolo A, Craig PS, Wassermann M, Takahashi K, de la Rue M. Ecology and Life Cycle Patterns of Echinococcus Species. Adv Parasitol. 2017;95:213–314.

Romig T, Omer RA, Zeyhle E, Hüttner M, Dinkel A, Siefert L, Elmahdi IE, Magambo J, Ocaido M, Menezes CN, Ahmed ME, Mbae C, Grobusch MP, Kern P. Echinococcosis in sub-Saharan Africa: emerging complexity. Vet Parasitol. 2011 Sep 8;181(1):43–7.

Sanger F, Nicklen S, Coulson AR. DNA sequencing with chain-terminating inhibitors. Proc Natl Acad Sci U S A. 1977 Dec;74(12):5463–7.

Sievers F, Wilm A, Dineen D, Gibson TJ, Karplus K, Li W, Lopez R, McWilliam H, Remmert M, Söding J, Thompson JD, Higgins DG. Fast, scalable generation of high-quality protein multiple sequence alignments using Clustal Omega. Mol Syst Biol. 2011 Oct 11;7:539.

Sneath PHA, Sokal RR. Numerical taxonomy. San Francisco (CA): W.H. Freeman; 1973.

Széll Z, Sréter-Lancz Z, Sréter T. Evaluation of faecal flotation methods followed by species-specific PCR for detection of Echinococcus multilocularis in the definitive hosts. Acta Parasitol. 2014 Jun;59(2):331–6.

Towner JS, Rollin PE, Bausch DG, Sanchez A, Crary SM, Vincent M, Lee WF, Spiropoulou CF, Ksiazek TG, Lukwiya M, Kaducu F, Downing R, Nichol ST. Rapid diagnosis of Ebola hemorrhagic fever by reverse transcription-PCR in an outbreak setting and assessment of patient viral load as a predictor of outcome. J Virol. 2004 Apr;78(8):4330–41.

Tsai IJ, Zarowiecki M, Holroyd N, Garciarrubio A, Sánchez-Flores A, Brooks KL, Tracey A, Bobes RJ, Fragoso G, Sciutto E, Aslett M, Beasley H, Bennett HM, Cai X,Camicia F, Clark R, Cucher M, De Silva N, Day TA, Deplazes P, Estrada K, Fernández C, Holland PWH, Hou J, Hu S, Huckvale T, Hung SS, Kamenetzky L, Keane JA, Kiss F, Koziol U, Lambert O, Liu K, Luo X, Luo Y, Macchiaroli N, Nichol S, Paps J, Parkinson J, Pouchkina-Stantcheva N, Riddiford N, Rosenzvit M, Salinas G, Wasmuth JD, Zamanian M, Zheng Y; Taenia solium Genome Consortium, Cai J, Soberón X, Olson PD, Laclette JP, Brehm K, Berriman M. The genomes of four tapeworm species reveal adaptations to parasitism. Nature. 2013 Apr 4;496(7443):57–63.

Untergasser A, Cutcutache I, Koressaar T, Ye J, Faircloth BC, Remm M, Rozen SG. Primer3--new capabilities and interfaces. Nucleic Acids Res. 2012 Aug;40(15):e115.

Veit P, Bilger B, Schad V, Schäfer J, Frank W, Lucius R. Influence of environmental factors on the infectivity of Echinococcus multilocularis eggs. Parasitology. 1995 Jan;110 (Pt 1):79–86.

Wachira TM, Bowles J, Zeyhle E, McManus DP. Molecular examination of the sympatry and distribution of sheep and camel strains of Echinococcus granulosus in Kenya. Am J Trop Med Hyg. 1993 Apr;48(4):473–9.

Waterhouse AM, Procter JB, Martin DM, Clamp M, Barton GJ. Jalview Version - a multiple sequence alignment editor and analysis workbench. Bioinformatics. 2009 May 1;25(9):1189–91.

Wiedenmann A, Krüger P, Dietz K, López-Pila JM, Szewzyk R, Botzenhart K. A randomized controlled trial assessing infectious disease risks from bathing in fresh recreational waters in relation to the concentration of Escherichia coli, intestinal enterococci, Clostridium perfringens, and somatic coliphages. Environ Health Perspect. 2006 Feb;114(2):228–36.

Ye J, Coulouris G, Zaretskaya I, Cutcutache I, Rozen S, Madden TL. Primer-BLAST: a tool to design target-specific primers for polymerase chain reaction. BMC Bioinformatics. 2012 Jun 18;13:134.

Zheng H, Zhang W, Zhang L, Zhang Z, Li J, Lu G, Zhu Y, Wang Y, Huang Y, Liu J, Kang H, Chen J, Wang L, Chen A, Yu S, Gao Z, Jin L, Gu W, Wang Z, Zhao L, Shi B, Wen H, Lin R, Jones MK, Brejova B, Vinar T, Zhao G, McManus DP, Chen Z, Zhou Y, Wang S. The genome of the hydatid tapeworm Echinococcus granulosus. Nat Genet. 2013 Oct;45(10):1168–75.

